# The impact of incidental anxiety on the neural signature of mentalizing

**DOI:** 10.1101/2023.07.25.550477

**Authors:** Li-Ang Chang, Jan B. Engelmann

## Abstract

While the effects of anxiety on various cognitive processes, including memory, attention, and learning, have been relatively well documented, the neurobiological effects of anxiety on social cognitive processes remain largely unknown. We address this gap using threat-of-shock to induce incidental anxiety while participants performed two false-belief tasks, a standard and an economic-games version. During belief formation and belief inferences, regions in a canonical social cognition network showed activation reflecting mentalizing, including the temporoparietal junction (TPJ), precuneus, and dorsomedial prefrontal cortex (dmPFC). At the same time, we found threat-related suppression of social cognition regions during belief inferences. A conjunction analysis confirmed that a network of regions was simultaneously *engaged* during mentalizing and *suppressed* by anxiety: bilateral TPJ, bilateral IFG, and putamen. We examined how threat impacted the connectivity between seed regions from the conjunction analyses and its targets. During belief formation, we found that anxiety suppressed the connectivity between the precuneus seed and two key mentalizing nodes, the dmPFC and right TPJ. Moreover, during belief inferences threat specificallty suppressed belief-based connectivity between putamen and its targets in IPS and dlPFC, and dispositional distress significantly modulated threat-related suppression of connectivity between the left TPJ seed and left IPS. Our results highlight important effects of incidental and dispositional anxiety on specific nodes of the social cognition network. Taken together, our study uncovers novel interactions between the reward, social cognition, and attentional systems, indicating that social cognitive processes rely on support from other large-scale networks, and that these network interactions are disrupted under incidental anxiety.

## Introduction

Because much of human everyday behavior occurs in the presence of some underlying background emotion, it is becoming increasingly important to understand how incidental emotions influence cognitive processes that support behavior. Anxiety is an emotion that is particularly prevalent for human behavior (Bandelow et al. 2015), as its presence signals the occurrence of negative events that can be crucial for survival. Recent research has begun to investigate the effects of such incidental anxiety on cognitive processes including attention (Bar-Haim et al., 2007; Bradley et al., 2010; Cisler & Koster, 2010; MacLeod & Mathews, 1988), memory (Andreotti et al., 2011; Balderston et al., 2017; Bolton & Robinson, 2017; Vytal et al., 2013), and learning (Becker et al., 2019; Berghorst et al., 2013; Browning et al., 2015; Cavanagh et al., 2019; DeVido et al., 2009; Grupe, 2017; Robinson et al., 2013; Safra et al., 2018; Stevens et al., 2014; Ting et al., 2020). Despite this recent surge in the interest concerning the effects of incidental anxiety on cognitive processes and its neural correlates, relatively little is known about its impact on arguably one of the most important cognitive processes for human interaction, namely social cognitions such as mentalizing. In the current paper we aim to address this significant gap in the literature and investigate how incidental anxiety impacts the underlying neural circuitry that supports mentalizing.

Mentalizing (also commonly referred to as Theory of Mind) is defined as the ability to attribute mental states to others, such as beliefs, intentions and desires, and thereby enables reasoning about others’ mental states (Frith & Frith, 2006; Premack & Woodruff, 1978). As such, mentalizing supports human interaction in important ways. Previous work has begun to delineate the effects of anxiety on social cognition. Clinical work suggests that specific anxiety disorders are associated with diminished mentalizing abilities. Compared to healthy controls, patients with social anxiety disorder (SAD) have been shown to perform poorer on the reading the mind in the eyes task (RMET) (Hezel & McNally, 2014), as well as other tasks including the Movie for the Assessment of Cognition (MASC) task (Washburn et al., 2016). Recently, a number of meta-analyses have further substantiated the link between pathological anxiety and a diminished ability to understand and reason about others’ perspective. Not only has social anxiety been found to negatively affect social cognitive capacity in children and adolescents (Pearcey et al., 2021), but this association also extends to other anxiety disorders and older patients. Sloover et al. (2022) observed a pattern of impaired mentalizing abilities across various patient groups, including those with anxiety disorders, obsessive-compulsive and related disorders, as well as trauma and stress-related disorders. In a more recent meta-analysis, Chevalier et al. (2023) focused on moderators of mentalization, including developmental components and sample characteristics, as well as the different measures of mentalizing and anxiety used across studies. Their results show a general and negative association between (unspecified, social and generalized) anxiety (as well as internalizing difficulties) and mentalizing performance across multiple tasks and questionnaires, while no moderating effects of sample type and developmental period were found. This suggests that anxiety is associated with mentalizing deficits in both clinical and subclinical samples and across developmental periods, and further substantiates a mild to moderate decrement in mentalization capacity that is prevalent across most anxiety disorder spectra.

The notion that anxiety suppresses social cognition is further supported by studies that induce incidental anxiety in healthy subjects, demonstrating causal effects of incidental anxiety that move beyond cross-sectional and correlational experimental designs. An initial set of experiments by Todd et al. (2015) induced anxiety via an autobiographical memory task before healthy participants completed a number of spatial and conceptual perspective taking tasks across multiple experiments. Results indicate that anxious participants were more likely to use their own spatial perspective in descriptions of object locations, showed more egocentric interference and assumed that others used knowledge only they had when inferring others’ beliefs. Jointly, these results indicate that under anxiety participants exhibited greater egocentrism compared to participants experiencing neutral and other negative emotions, such as anger and disgust.

Despite these initial behavioral findings on the effects of anxiety on mentalizing, relatively little is known about how incidental anxiety affects the underlying neurobiology that supports social cognition (see von Dawans et al., 2021 for an overview of research in the related field on stress). The neurobiological basis of both mentalizing and anxiety have been delineated separately by extensive prior neuroimaging work in social and affective neuroscience. Several meta-analyses agree on a core neural network that supports mentalizing across different tasks, consisting of bilateral temporoparietal junction (TPJ), medial prefrontal cortex (mPFC), superior temporal sulcus (STS), temporal pole (TP) and precuneus (sometimes including posterior cingulate cortex, PCC) (Amodio & Frith, 2006; Bzdok et al., 2012; Decety & Lamm, 2007; Mar, 2011; Mitchell, 2009; Molenberghs et al., 2016; Schurz et al., 2014; Van Overwalle, 2009). Paralelly, a recent fMRI meta-analysis that summarized the past two decades of neuroimaging research on anxiety identified a “core anxiety” network whose regions include the amygdala, the bed nucleus of the stria terminalis (BST), periaqueductal gray, midcingulate cortex, and anterior insula (Chavanne & Robinson, 2021). Of importance for the current study is the finding that these regions are involved in *both* clinical anxiety and pre-clinical studies that induce anxiety, suggesting that emotional challenges recruit regions within this network regardless of pre-existing symptomatology related to anxiety (Chavanne & Robinson, 2021).

While relatively little is known about the neurobiological effects of incidental anxiety on mentalizing, a prior neuroimaging study investigated the impact of induced stress on participant’s performance in the reading the mind in the eyes test (RMET). The authors compared two different stress induction approaches, general stress induction (GSI) and attachment-related stress induction (ASI) that asked subjects to think about stressful circumstances relevant to interpersonal relationships (Nolte et al., 2013). The fMRI results showed that attachment-related stress induction suppressed several mentalizing areas as compared with the non-social stress induction condition, particularly in the left posterior superior temporal sulcus (STS), left temporoparietal junction (LTPJ) and left inferior frontal gyrus (IFG). While the previous regions (STS/TPJ) are part of a core social cognition network, the IFG is commonly considered a cognitive control region, that can also be viewed as part of an extended social cognition network (Arioli et al., 2021).

Understanding the perspectives of others, which requires distinguishing between self and others and inhibiting one’s own perspective, inherently involves cognitive and inhibitory control areas such as the inferior frontal gyrus (IFG) (Lieberman, 2007). The role of this neural substrate in social cognition might be evidenced in conditions such as Autism Spectrum Disorder (ASD), where children exhibit diminished IFG activation while observing and imitating emotional expressions (Dapretto et al., 2005). Similarly, via employing a whole-brain voxel-based lesion symptom mapping approach, Dal Monte et al. (2014) found that lesions in the left IFG correlate with poorer RMET performance in traumatic brain injury patients. More recently, Engelmann et al. (2019) identified VLPFC as part of a network of regions whose connectivity strength with TPJ is specifically associated with trust game investments, and, at the same time, showed that anxiety induced by unpredictable threat of shock, significantly suppressed the involvement of the VLPFC in trust and non-social investment decisions. Jointly, the evidence suggests therefore that anxiety can suppress the activation of neural circuitry that supports social cognition.

On the other hand, patients with SAD show hyperactivation in brain regions associated with mentalizing, compared to healthy controls during multiple tasks, including working memory tasks using faces with self-referential evaluative comments (Yoon et al., 2016) and a self-referential emotion regulation task that involved viewing tasks of aversive compared to neutral images (Gaebler et al., 2014). Such hyperactivation, occurring especially in midline structures such as the mPFC and PCC, is particularly associated with a higher degree of individual negative affectivity (Lemogne et al., 2011). For instance, Rabany et al. (2017) show that individuals with anxiety disorders exhibit enhanced resting-state connectivity between mPFC and PCC. At the same time, some studies suggest that hyperactivation found in SAD patients is not limited to midline structures, but can also involve the TPJ (Boehme et al., 2015; Gaebler et al., 2014; Yoon et al., 2016). Such hyperactivation within the mentalizing network during self-referential processing in patients with SAD may reflect enhanced effort to focus on and understand others as a compensatory response to increased egocentricity that is likely associated with a diminished mentalizing capacity (e.g., Silani et al., 2013). Moreover, it is worth noting that these studies primarily focus on dispositional anxiety and anxiety disorders. Thus, it remains unclear whether incidental anxiety, which arises from environmental triggers, affects the mentalizing network in the same way. Results of Chavanne & Robinson (2021) showing significant overlap in anxiety-related activation in clinical samples and after anxiety induction in healthy controls tentatively suggest similar neural effects of clinical and induced anxiety. We test here whether this extends to the neural correlates of the effects of incidental anxiety on social cognition.

Social cognitive processes also crucially support social interaction during economic games. If anxiety interferes with these social cognitive processes during economic game interaction, it should lead to distorted economic behavior. Feldman Hall et al. (2015) applied the cold pressor test (CPT) to induce stress before participants performed a trust game. Results showed diminished trust compared to a non-social control condition, specifically after stress induction. These results were confirmed and extended by a more recent fMRI experiment in which participants played a trust game under conditions of incidental anxiety, induced via threat-of-shock (Engelmann et al. 2019). Behavioral results confirmed the earlier findings, showing decreased levels of trust under conditions of incidental anxiety compared to safety. At the neural level, the authors demonstrated supressed left TPJ activity and functional coupling between left TPJ and amygdala under conditions of anxiety. Moreover, while the fucntional connectivity between the left TPJ and its targets in pSTS (posterior superior temporal sulcus), dmPFC, amd vmPFC were associated with the transferred amount in the trust game (but not in the non-social control condition), anxiety speficifcally disrupted this brain-behaviour relationship between left TPJ and pSTS. Jointly, these results indicate that interpersonal behaviors, such as trust, are disrupted by anxiety. Moreover, the finding that the disruption of this behavior correlates with specific suppression of activity and connectivity between regions involved in mentalizing further suggests that these effects may be driven by a disruption of social cognitive processes.

In the current study, we aim to provide direct evidence for the disruption of mentalizing in the presence of incidental anxiety and delineate the effects of anxiety on the activity and connectivity within the social cognition network. Specifically, we ask whether and how anxiety influences the ability to mentalize and to what degree anxiety disrupts the social cognition network that supports mentalizing. To accomplish this, we asked our subjects to complete two different false-belief tasks for which they first read vignettes about life stories (the standard FBT), or about interactions in economic games (the economic-game FBT). Our participants did this either in the presence of threat of shock, or in its absence. We analyze the effect of threat of shock on mentalizing-related activation and connectivity during two periods: (1) a vignette reading period, in which participants were required to form beliefs about agents in vignettes, and (2) during a question period, in which participants were asked to answer incentivized questions about the agents’ beliefs and intentions in the belief condition and about simple outcomes and payment distributions in the outcome condition. Based on results in prior research, we expected suppressed activity and connectivity during threat in mentalizing regions. Given our previous results (Engelmann et al., 2019), we expected such threat-related suppression particularly in the TPJ.

## Materials and Methods

### Subjects

39 volunteers participated in the current experiment (18 males, aged 18 - 33, mean (SD) = 22.51 (4.03) years). Two subjects were excluded from data analyses, one due to excessive head motion (> 2 voxels (6 mm) in translation and rotation) and one due to subthreshold task performance (mean accuracy < 3 standard deviation of the sample mean). The final dataset for the main analysis therefore included 37 subjects. For analyses of individual differences one further subject needed to be excluded due to technical issues with the online questionnaire that led to data loss. Participants were recruited from the subject pool of the Behavioral Science Lab at the University of Amsterdam (https://www.lab.uva.nl/lab/home) and provided informed consent to procedures that were approved by the Ethics Review Board of the Faculty of Social and Behavioral Sciences of the University of Amsterdam.

### Timeline of procedures

In the first part of the experiment, participants were invited to fill in an online prescreening questionnaire and a battery of personality measures via Qualtrics. The questionnaire battery was adapted from (Engelmann et al., 2019; Engelmann et al., 2020) and is outlined in detail below. Subjects received 14 Euros for completing this part of the study. In the second part of the experiment, subjects were invited to the Behavioral Science Lab at the University of Amsterdam. They were asked to read the instructions thoroughly, after which their questions were discussed with the experimenter, followed by a quiz that assessed their understanding of the task, and, in particular, the economic games (Trust and Ultimatum Games). Participants then completed 12 practice trials outside the scanner, which were repeated until performance was at least 66% (only 3 subjects required repetition and all of them passed the criteria on their second try). After entering the scanner, participants first underwent shock calibration and were then familiarized with the MRI compatible button box. To control for possible (de)sensitization to the electric shocks, a second shock calibration was conducted at the halfway point of the experiment (after completion of run 2 out of 4). After scanning, subjects filled out an exit questionnaire which included additional manipulation checks and personality measurements (Honesty-Humility scale of the HEXACO-PI-R (Lee & Ashton, 2010) and the Dark Factor of personality (Moshagen et al., 2018). Subjects were paid an average of 32.32 Euros for their participation via the online reimbursement system of the University of Amsterdam.

### False-belief task description

We employed a vignette-based false belief task to investigate the role of anxiety in social cognition, focusing in particular on belief formation, which occurred during the vignette period, and belief inferences, which occurred during the question period. Specifically, we combined well-established and slightly modified (to improve readability) vignettes from prior research (adapted from Bruneau et al., 2012; Saxe & Kanwisher, 2003), with a set of novel vignettes that outline interactions in well-established economic games. For a detailed analysis of these stimuli and their neural correlates, see our previous paper (Chang et al., 2023). To complete the task, participants were asked to first read the vignettes and subsequently answer incentivized questions about the beliefs of protagonists and outcomes in different experimental conditions.

As shown in Error! Reference source not found.**A,** the experiment employed a within-subjects factorial design with the factors Domain (economic games, life stories), Belief (belief, outcome), and Threat (threat, safety). This design allowed us to test the effects of anxiety, manipulated via the well-established threat of shock procedure (see Threat of Shock procedure), on theory of mind related neural activations (reflected via the contrast belief - outcome) in economic games and life stories. To give an example of the stimuli, in the Life Story-Outcome condition, the subjects read about events that happen to another person. They were asked to answer questions about an objective description of the consequence of the event. In the Life Story-Belief condition, the subjects were asked to infer the beliefs or intentions of the protagonist in the scenario. Economic game stimuli, on the other hand, were based on hypothetical interactions between different partners in well-established economic games, including the trust game (TG, Engelmann, 2010) and the Ultimatum game (UG, Thaler, 1988), and therefore required a well-founded understanding of the game setup and payout conditions to correctly answer the question. In the Economic Game-Outcome condition, the subjects were asked to calculate the payoff for one of the interaction partners based on the rules of the economic game that were outlined in detail and subsequently quizzed during the instructions. In the Economic Game-Belief condition, the subjects were required to infer the (false) beliefs of one of the interaction partners during either a Trust or Ultimatum Game interaction. The following provides an example of a trust game vignette in which participants were required to infer the false belief of one of the interaction partners:

> Eline and Kostas play a Trust Game. Each of them gets 15 Euros. Eline is the investor and sends all her money to Kostas. Kostas is confused and thinks that now they both have 60 Euros, so he sends back nothing. Eline hides her face in her hands. Question: Eline probably thinks that Kostas: is greedy (correct) / is confused (incorrect).

Answers were incentivized at a piece-rate of 20 cents for each correct answer, leading to additional earnings of up to €19.20 for answering all 96 questions correctly. Final payment for participation in this experiment was thus a maximum of 33.20 Euros (€19.20 for correct answers plus €14 for questionnaire completion). It was stressed in the instructions that answering correctly and on time on each trial was in the participants’ best interest in terms of the monetary payoff, but that task performance cannot influence the number (or intensity) of the electric shocks.

### Experimental Design and Trial Timing

We used a hybrid fMRI design that included a blocked component, namely the presence and absence of anxiety induced via Threat of Shock, and event-related components, namely belief formation during the vignette and belief inferences during the question periods. A specific advantage of this fMRI design is that it allows us to induce anxiety for a prolonged period of time. Furthermore, this approach reduces task switching demands (see Engelmann et al., 2015; Engelmann et al., 2019; Ting et al., 2020). At the beginning of each block, participants were informed of the current condition via a block cue that indicated whether vignettes were presented in the economic games or life stories domain, whether questions would assess beliefs or outcomes, and whether shocks would be administered at unpredictable time points throughout the next block or not. The Threat condition was also communicated to participants via a constant background color that remained the same for the period of each block, such that for about half the participants a green background color was associated with threat blocks, while a blue background color was associated with safety. For the remaining subjects, this association was reversed. Participants learned about the color-threat association in the instructions and during the practice task outside the scanner. **Figure 1A** illustrates the sequence and timing of an example trial. Each block started with a block cue that was shown for 3000ms and provided three pieces of information to the subjects, namely Vignette and Belief Domain, and the current Threat condition. The example in the figure shows an Econ-Belief-Shock condition, indicating that the 3 vignettes in the current block concern economic game scenarios, and subjects need to focus on inferring character’s beliefs, with the possibility of receiving an unpredictable number of shocks at unpredictable time points throughout the block. The block cue is then followed by another blank screen containing a fixation cross presented for a jittered duration (mean = 4000ms). Thereafter, subjects were asked to read a vignette, for which they were given 10000ms. The vignette display was followed by a question period which gave participants a maximum of 7000ms to answer the question by choosing from two options, one correct and one incorrect option. The position of the correct option was randomized across trials. To ensure that subjects paid attention continuously throughout the experiment, correct answers within the 7000ms limit yielded a piece-rate payment of 20 cents that was added to participants’ cumulative earnings. The question period terminated when participants provided their answer via button press, or after the maximum allotted time of 7000ms and was followed by feedback about performance (shown for 500ms plus the remaining time from the question period, i.e., 7000ms – RT, see **Figure 1A**) indicating whether participants were correct, wrong, or answered too slowly. Note that our procedure prevented participants from accelerating progress on the task by responding faster, because it ensured that threat and safe blocks maintained the same average length. Participants were made aware in the instructions that they could not speed up responding in threat blocks to avoid being shocked, and that their responses on the task in no way influenced the amount, intensity, and timing of electrical shocks. Each trial was followed by an intertrial interval showing a fixation cross for a jittered period (mean = 4000 ms). The experiment consisted of a total of 96 trials distributed across 4 runs. Each run contained 8 blocks, within which 3 trials were presented in the same condition.

**Figure 1.**
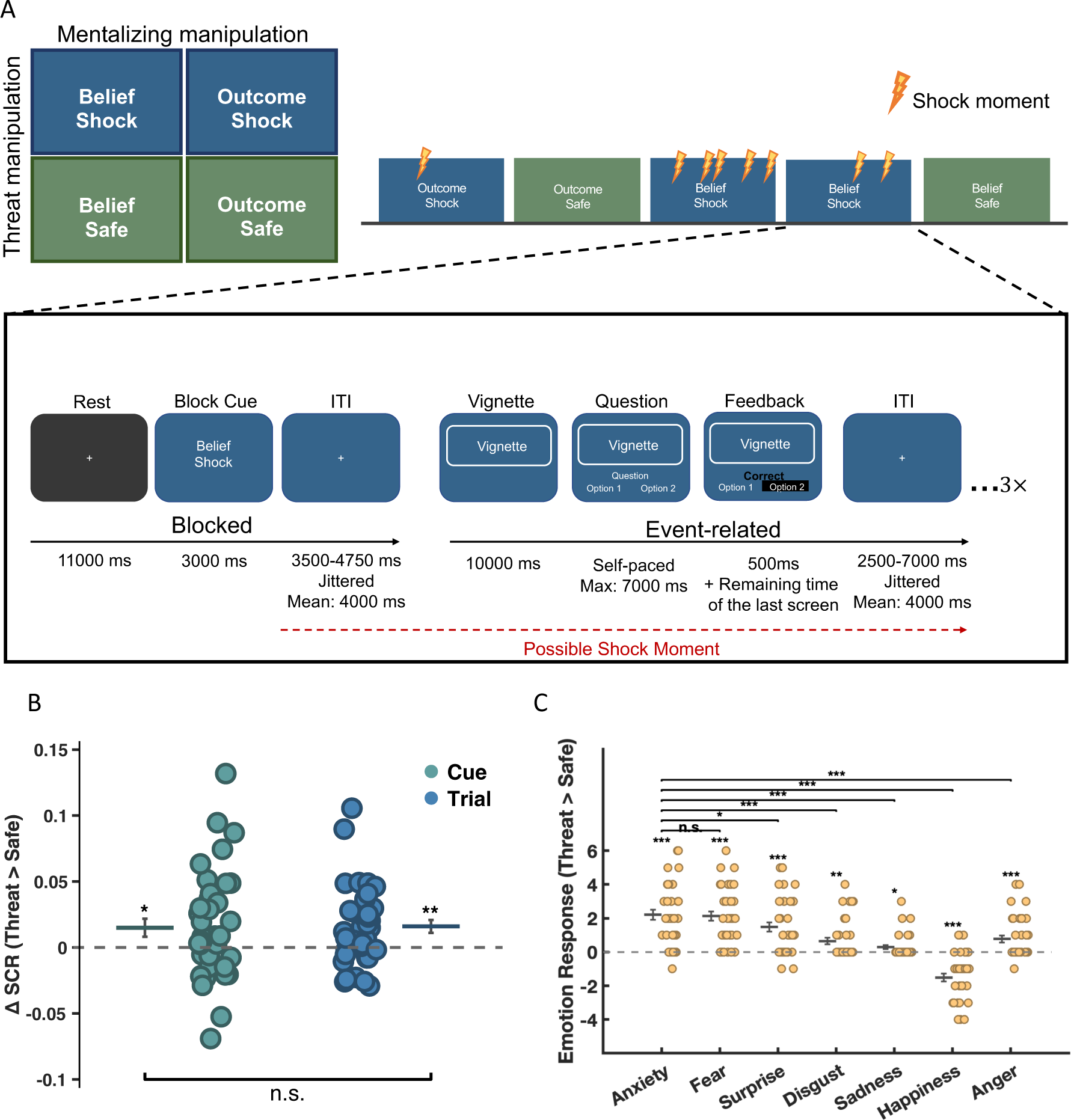
Experimental conditions, trial sequence and timing and manipulation checks. A) 2×2 factorial design of the mentalizing and threat manipulations. On right, implementation of the hybrid fMRI design in which the Threat and Belief factors were consistent for the duration of a block, as indicated by the block cue, and vignettes, questions and feedback were presented in an event-related fashion. Feedback gave information on correct, incorrect or slow responses, where only the former was paid at a 20c piece rate. The example shows the Econ-Belief-Threat condition. (Note: the background color associated with the shock condition was counterbalanced across subjects, but remained the same for each subject from practice until the end of the experiment). Manipulation checks show significant threat effects on (B) skin conductance responses and (C) self-reported emotion. (B) Significantly larger SCR responses were observed in the threat compared to the safe condition during two phases of the experiment: the cue period (green dots) and throughout the trial period (blue dots). Each dot in the figure represents individual’s estimated parameter extracted from the contrast of (threat > safe) either during the cue period or the trial period after cue offset in the general linear model (GLM). (C) Subjects reported significantly higher levels of fear, anxiety and surprise in threat compared to safe blocks, while reporting greater happiness in safe compared to threat blocks. In addition, the increase in self-reported anxiety in threat relative to safe blocks is significantly greater compared to all other emotions, except for fear. In both subplots (B and C), the horizontal line represents the mean, and the black error bar represents the standard error of the mean (SEM). Each dot represents an individual participant p < .1 ∼ ; p < .05 *; p <.01**; p < .001***

### Anxiety Induction using Threat of Shock

Anxiety was induced via the well-established threat-of-shock procedure (Engelmann et al., 2015; Engelmann et al., 2019; Grillon et al., 2004; Ting et al., 2020), which administers unpredictable and mildly painful electric shocks in the Threat condition. The carpometacarpal joint of the participants’ non-dominant hand (always the left hand in the current study) was connected to a DS5 Isolated Bipolar Constant Current Stimulator (Digitimer Ltd.) through MRI compatible electrodes and wires. A DS5 stimulator generated stable electric shocks with a fixed maximum input of 5V, maximum output current of 25mA, and a fixed shock duration of 50ms across all subjects. The specific intensity of the electric shock was customized to each subject’s pain intensity threshold using a well-established and automated calibration procedure (e.g., Ting et al., 2021). During calibration, subjects were instructed to press a button to trigger the electrical shock themselves. After experiencing the shock, they were asked to rate how painful their experience of the electric shock was via a visual analogue scale (VAS) that ranged from 0 (no pain) to 10 (extreme pain). The automated procedure used a staircase algorithm in which ratings below 7 led to an increase of the shock amplitude by 10% of the maximum output intensity (5V, 25mA, duration 50ms), ratings between 7 and 9 led to no change, while ratings of 10 led to a 10% intensity decrease. The calibration procedure terminated once three consecutive ratings of 7 to 9 were provided by each participant. If the subject provided only ratings below 7, the calibration would terminate after reaching the maximum output intensity (this occurred in 7 participants after the first calibration). If participants answers were highly inconsistent, the calibration terminated after 20 trials and the value of the final trial was used as shock amplitude (this did not occur for any participants). As noted in the timeline of the procedure section, the first calibration was conducted before the first run, and the second calibration was conducted at the halfway point of the fMRI experiment (after completion of the second run) to control for (de)sensitization effects. The average stimulation intensity selected by participants during the first calibration was 52.16% of the maximum output intensity, which increased for the second calibration to 61.35 %. This indicates that participants selected a stimulation intensity that was significantly above the starting value of 10% (t_36_ = 9.16, *p* = 6.15×10^-11^), and that this value further increased for the second half of the experiment (t_36_ = 4.62, *p* = 4.81×10^-5^, **Figure S1**). Jointly, these results provide evidence suggesting that our procedures worked well, as participants did not choose to receive the lowest possible stimulation amplitude throughout the experiment. Moreover, we also find desensitization reflected by a significant increase in average stimulation intensities after the second calibration. This indicates that participants requested even higher stimulation amplitudes during the second calibration, and moreover, that our procedures can account for such desensitization effects.

Electrical stimulation procedures throughout the fMRI (and pilot) experiments were adapted from our prior studies (Engelmann et al., 2015; Engelmann et al., 2019; Ting et al., 2020). Specifically, to maximize unpredictability of stimulation time points to subjects and thus to generate a robust anticipatory anxiety effect (Schmitz & Grillon, 2012), both the number of shocks and the time points of electric shocks were pseudo-randomized for each threat block. Electrical shocks could be administered any time (but were kept at least 1000ms apart) in the period spanning from the offset of the block cue until completion of the last trial in a given block. Possible time points of shocks were drawn from a random normal distribution throughout this time window. The number of shocks per block in each condition were pseudo-randomized for each condition within each run by selecting from a gamma function (with the values 1, 2, 2, 5) via random draw without replacement. This ensured that participants experienced the same number of shocks in each Threat condition (across the different Domain and Belief conditions). Finally, neither participants’ response speed nor their accuracy on the task could influence the intensity or the frequency of getting shocked, which we explicitly told our participants in the instructions. Jointly, these procedures, including the threat block cue, the unpredictable amount and time points of shock administration and the sustained period throughout which electrical stimulation was possible, have successfully induced prolonged anxiety in our previous studies (Engelmann et al., 2015, 2019; Ting et al., 2021).

### SCR recording and analysis approach

To ensure the success of the emotional manipulation we measured participants’ skin conductance response (SCR). SCR data were measured using the BrainAmp ExG MR (Brain Vision, Morrisville, NC, USA) with two MR-compatible finger electrodes attached to the index and middle fingers of the participants’ nondominant left hand after conductance gel was applied. SCR data were sampled at 5000 Hz and SCR recordings were done on a per-run basis to reduce low frequency drift.

SCR data were preprocessed and analyzed using PsPM (PsychoPhysiological Modelling, Bach et al., 2009). All signals were filtered using a bidirectional Butterworth band pass filter with cutoff frequencies of 0.0159 Hz and 5 Hz and down-sampled from 5000 Hz to 10 Hz and subsequently z-transformed. A general linear model (GLM) was performed to estimate the ToS effect (threat vs. safe) during block cue presentation, as well as during the trial period. Our GLM for SCR analysis included a total of five event-related regressors, namely shock cue, safe cue, shock trial, safe trial, and the actual shock moments. All events were convolved by a canonical SCR function with a time derivative (Bach, 2014). Importantly, we also added a regressor in the model to take the shock moment *per se* into account, which enabled us to capture the affective component of the SCR signal during threat and shock trials, which is prolonged and independent of the actual shocks that could occur at any time point throughout a given trial.

### Functional magnetic resonance imaging

#### FMRI data acquisition

FMRI data were collected using a 3.0 Tesla Philips Achieva TX MRI scanner (Philips Medical Systems, Best, The Netherlands) using a 32-channel head coil. T1-weighted structural images were acquired (1 × 1 × 1 mm voxel size resolution of 220 slices, slice encoding direction: FH axial ascending, without the slice gap, TR = 8.2 ms, TE = 3.7 ms. flip angle = 8°). Functional images were acquired using a T2*-weighted gradient-echo, echo-planar pulse sequence (3.0 mm slice thickness, interslice gap = 0.3 mm, 36 ascending slices, TR = 2000 ms, TE = 28 ms, Flip angle = 76.1°, and with 240 mm field of view). In addition, to correct EPIs for signal distortion, we also conducted an additional field-map scan at the half-way point of the experiment. We implemented a Phase-difference (B0) scan with reference (2.0 × 2.0 × 2.0 mm voxel size resolution, axial ascending direction, without slice gap, TR = 11 ms, TE = 3 ms, flip angle = 8°).

#### FMRI preprocessing and general linear model (GLM) analyses

FMRI data preprocessing and analysis was carried out with SPM12 (Wellcome Department of Cognitive Neurology, London, UK). Preprocessing followed the following steps: First, all functional images were simultaneously realigned to the first volume of the first run using septic b-spline interpolation and unwarped (using B0 maps) using the realign and unwarp function in SPM, followed by slice timing correction. Afterward, T1-weighted structural images were co-registered with the functional images and then segmented into six different tissue classes using the segment function in SPM12. Next, all images were normalized to the Montreal Neurological Institute (MNI) T1 template with the forward deformation parameters obtained from segmentation. Lastly, all functional images were smoothed using spatial convolution with Gaussian kernel of 6 mm at full width half maximum (FWHM).

Statistical analyses were carried out using the general linear model (GLM). Our statistical model reflects the factorial design of the current experiment by including separate regressors for each condition of the factor Threat (safe, threat) and Belief (false belief vs. outcome), separately for both the vignette and question periods. All regressors were modeled using a canonical hemodynamic response function (HRF). To best capture mentalizing during the question period, we used a variable epoch model from the onset of the question until option choice (button press). We also modeled regressors of no interest, which include each block cue, the feedback period, shock moments and the Domain condition (economic games vs. life story), note that we show in a separate paper that economic games and life story domains recruit similar social cognition regions during both periods (see Chang et al., 2023), which is why we include this factor here as a regressor of no interest), as well as omitted trials during which no response was made by the subject. While omissions were rare (on average 0.55%), these trials were modeled explicitly to ensure that we only included trials on which we are certain participants paid attention to the task. In addition, the six motion parameters derived from the realignment procedure were modeled as regressors of no interest.

Aim of the current study was to investigate the impact of anxiety induced via ToS on the neural correlates of mentalizing. To this end, the main focus of the analysis was the main effect reflecting the engagement of social cognition regions to infer the (false) beliefs of the vignette’s protagonists (Belief > Outcome) and the effect of Threat on these neural activations (Safe > Shock; Shock > Safe). To examine whether threat impacted on the neural network involved in theory of mind, we conducted a conjunction analysis, which identified voxels that showed both, a social cognition effect, and at the same time an effect of threat. We conducted the conjunction analyses separately for the vignette period, reflecting social cognitive processes related to belief formation, and during the question period, reflecting social cognitive processes related to belief-based inferences. Individual contrast maps that were entered into conjunction analyses were FWE-corrected at cluster level with a cluster-forming threshold of *p* < 0.001. Conjunction analyses were conducted under the conjunction null hypothesis, requiring that all comparisons in the conjunction are individually significant (Nichols et al., 2005). Follow-up functional connectivity analyses were then conducted on those regions that are engaged during false belief formation and inferences and that show a simultaneous effect of Threat. To this end, we performed general psychophysiological interaction (gPPI) analyses to assess the more extended circuitry that supports social cognition, and how this extended network is impacted by Threat.

#### Connectivity Analyses: Generalized Psychophysiological interaction (gPPI) Analyses

To assess how threat impacts effective connectivity during mentalizing, we conducted generalized psychophysiological interactions (gPPI) using the CONN toolbox (www.nitrc.org/projects/conn) (Whitfield-Gabrieli & Nieto-Castanon, 2012). To increase sensitivity and specificity for functional connectivity analyses, we again preprocessed the data in the CONN toolbox using the indirect segmentation and normalization pipeline in CONN, which is largely equivalent to our preprocessing steps above, but included the additional step of identifying and removing outlier scans from the analysis (ART, Whitfield Gabrieli). In accordance with the anatomical component-based noise correction method (Behzadi et al., 2007; Muschelli et al., 2014), additional denoising was conducted before functional connectivity analyses, which included 10 cerebrospinal fluid (CSF) and 10 white matter principal components as nuisance covariates, as well as motion parameters, their first temporal derivatives and quadratic extensions (total 24 parameters, (Friston et al., 1996), all scrubbed datapoints from the artefact removal step and all regressors and temporal derivatives. Low-frequency fluctuations were isolated using a low-pass temporal filter (.008 Hz) after denoising.

Seed regions for functional connectivity analyses were extracted from the conjunction maps identifying overlap between the threat manipulation (safe > threat) and mentalizing (belief > outcome) during the question period. These included bilateral TPJ (Left: -60, -61, 20, k = 83; Right: 60, -58, 35, k = 22), bilateral IFG (Left: -54, 23, 5, k = 29, Right: 51, 26, -7, k = 28), as well as left putamen (-28, 8, 5, k = 8). As no threat suppression area was identified during the vignette period, this conjunction analysis is done only for the question period.

## Results

### Threat of shock manipulation checks

#### Manipulation Check 1: Skin Conductance Responses

We first tested the effectiveness of our threat of shock procedure to induce anxiety by assessing our participants’ autonomic arousal across the threat and safe blocks. Our results demonstrate a robust and larger SCR signal at the moment of receiving the shock (t_36_ = 10.67, *p* <.001) compared to baseline. We observed a similar effect when the block cue was presented, with significantly increased SCR responses during threat relative to safety cues (t_36_ = 2.20, *p* = 0.03). Finally, increased SCR responses were observed throughout the entire trial period, that is from onset of the vignette until the end of the last trial, for threat relative to no-threat trials (t_36_ = 1.78, *p* =.002). Jointly, these results indicate that our threat of shock procedure induced increased autonomic arousal, both in terms of anticipatory affect that was triggered during the Shock Cue, and the incidental uncertainty of being under the threat of shock throughout shock trials (**Figure 1B**).

#### Manipulation Check 2: Self-reported Emotion

Next, we assessed the specific emotion associated with the increased arousal observed during threat blocks. Paired t-tests revealed that self-reported emotions differed significantly between threat and safe blocks. Specifically, subjects reported higher aversive emotion during threat compared to safe blocks, including Anxiety (t_36_ = 7.57, *p* = 6.02 × 10^-9^, d = 1.24); Fear (t_36_ = 7.94, *p* = 2.00 × 10^-9^, d = 1.31); Disgust (t_36_ = 3.40, *p* = 1.65 ×10^-3^, d = 0.56); Sadness (t_36_ = 2.58, *p* = 0.01, d = 0.42); and Anger (t_36_ = 3.81, *p* = 5.18 × 10^-4^, d = 0.63). Moreover, during threat blocks they felt more surprised (t_36_ = 5.56, *p* = 2.70 × 10^-6^, d = 0.91) and less happy (t_36_ = -6.45, *p* = 1.72 × 10^-7^, d = -1.06). We also tested whether threat of shock specifically increased self-reported anxiety to a larger extent than other aversive emotions. We find that anxiety was reported to be more intense compared to all other aversive emotions, except fear: Anger, t_36_ = 4.53, *p* = 6.22 × 10^-5^, d = 0.75; Disgust, t_36_ = 5.04, *p* = 1.35 × 10^-5^, d = 0.83; Sadness, t_36_ = 6.60, *p* = 1.12 × 10^-7^, d = 1.08; Surprise, t_36_ = 2.15, *p* = 0.04, d = 0.35; Fear (t_36_ = 0.29, *p* = 0.78, d = 0.05). These results indicate that our paradigm successfully increased anxiety, and did so specifically compared to other aversive emotions (**Figure 1B and C**).

In addition to significant psychophysiological and affective reactivity to shock administration, we also observe significant effects of electrical shock at the neural level. Specifically, administration of electrical shock, reflected by a regressor modeling the exact time point of electrical shocks, led to significant activation within the pain matrix, including the insula, anterior cingulate cortex, and the right somatosensory cortex (see **Figure S2**). Note that our fMRI models included a regressor for all shock time points to control for its effects.

#### Behavioral results

Goal of the behavioral analysis was to identify the effects of anxiety on performance and response times in the false belief task (FBT). To analyze the choice data, we conducted logistic regressions on trial-by-trial data that were implemented in the context of a generalized linear mixed-effects model (GLME). Two separate models included (1) responses on each trial (correct/incorrect) and (2) log reaction time as dependent variables, as well as Threat and Belief condition as fixed effects predictor variables (see **Tables S1** and **S2** for regression equations with all fixed and random effects). Task Domain, i.e. whether participants performed the standard or economic games FBT, was entered as regressor of no interest in the current analyses and is discussed in more detail in a prior analysis of this data set (Chang et al., 2023). Models were estimated via the mixed function of the AFEX package in R (Singmann et al., 2016) that relies on the lme4 package. We report results from models with the maximum possible random-effects structure (Barr, 2013). For reaction times, linear regressions using a full model structure with random slopes for the Belief factor, in addition to subjectwise random intercepts were employed. For accuracy, logistic regressions were employed. Including all random slopes led to overfitting, requiring us to remove random slopes such that all final models include a subjectwise random intercept only.

Results across different model specification show a significant main effect of Belief (*p* < 0.001) and a marginal effect of threat (*p* = 0.124, note: only in some model specifications, see **Table S1**). The effect of Belief indicates better performance in the Belief condition (mean percent correct = 0.98) compared to the Outcome condition (mean percent correct = 0.96). We show in a separate paper analysing different conditions of the same dataset (Chang et al., 2023) that this is largely driven by the outcome condition in the economic games, which is slightly more difficult as subjects were asked to compute the payouts of participants interacting in trust and ultimatum games. Inclusion of this control condition in our task is important, however, as it allows us to continuously check our subjects’ understanding of the payout structure of the economic games. Note that while significant, this effect is relatively small and reflects only a 2% improvement in the Belief compared to the Outcome condition. The marginal effect of threat (*p* = 0.124) in some model specifications indicates trend-level improvement in performance in the threat condition (mean percent correct = 0.975) compared to the safe condition (mean percent correct = 0.968). Note, however, that this effect does not persist across model specifications.

Response time results only showed a significant main effect for Belief (*p* = 0.035), but no main or interaction effects with the factor Threat (**Table S1**). The main effect indicates that in the belief condition response times were slower compared to the outcome condition (mean log RT in belief = 0.882; mean log RT in outcome = 0.844). Together with the accuracy results, these findings suggest that participants in the belief condition are both better and slower, hinting at potential speed-accuracy trade-offs. To test for the possibility that speed-accuracy trade-offs distort our results, we computed the Balanced Integration Score that corrects for this possibility (BIS, Liesefeld & Janczyk, 2019) for each subject and in each condition. Results from this analysis are largely comparable with what we report in mixed models above, but we find a slightly stronger trend effect for Threat reflecting non-significantly better performance in the threat condition (*p* = 0.107, **Table S2**) and further confirming this trend from the trial-by-trial analysis. The behavioral results of individual participants in each condition are shown in **Figure 3** below and reflect the mean performance of each participant across all trials within each condition.

**Figure 3.**
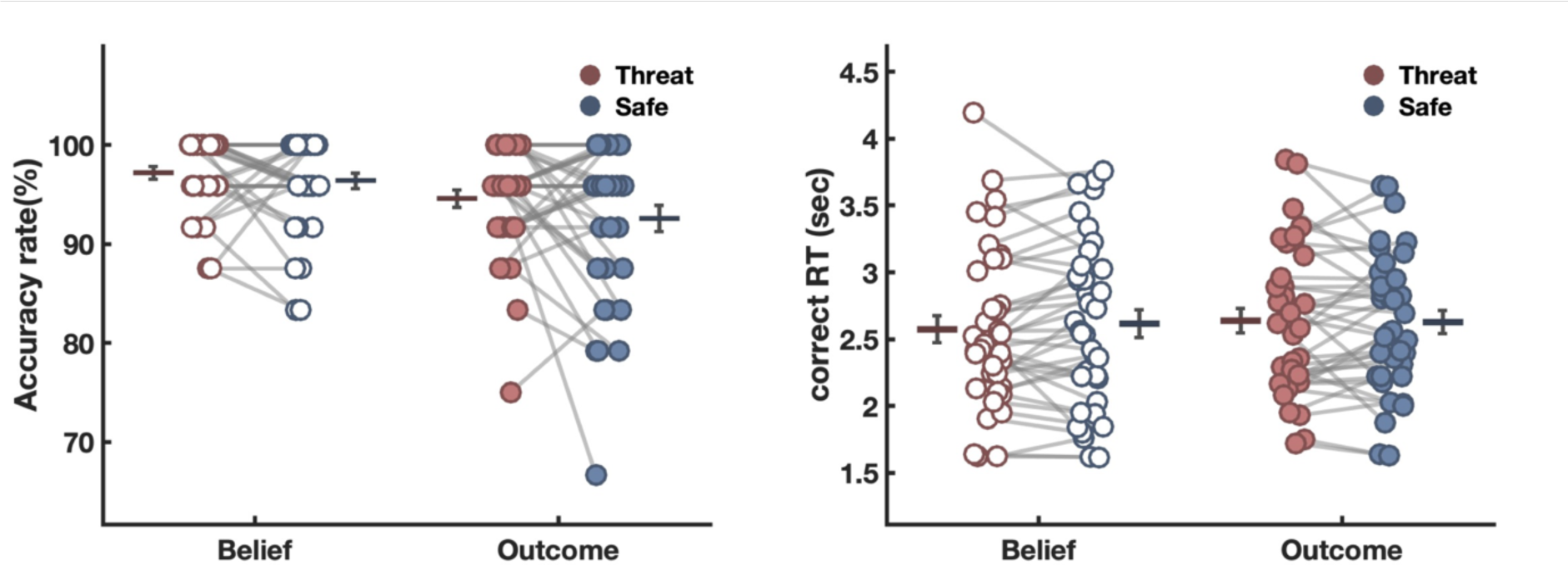
Mean of accuracy and correct RT in each condition. Average accuracy (left) and response time (right) for correct trials across Belief and Threat conditions. The horizontally line represents the mean, and the black error bar represents the standard error of the mean (SEM) in each condition. Each dot represents an individual participant, and the dots with gray connections indicate that they are from the same subject.

### FMRI results

#### Mentalizing-related activity during vignette and question periods

During the vignette period, we observed a main effect of belief formation (contrast: belief vs. outcome during vignette period) in the canonical social cognition network that includes bilateral temporoparietal junction (TPJ) (left: -48, -61, 26, k = 497; 48, right: -55, 23, k = 380), dorsomedial PFC (-6, 47, 35, k = 422), right superior temporal gyrus (48, -28, -1, k = 554), and precuneus (-6, -55, 29, k = 125) (**Figure S3A**) (**Table 1**). Belief-based activation during the question period was very similar and included many of the same regions (**Figure S3B**) (**Table 2**), but activation extent was more extensive. When subjects made belief inferences during the question period (contrast: belief vs. outcome during question period), we observed activity in left TPJ (extending to TP; -57, -28, -1, k = 3582), dmPFC (-9, 59, 32, k = 1343), precuneus (-3. -55, 29, k = 247), sensorimotor area (-48, -4, 50, k =94) as well as cerebellum (24, -73, -37, k =122; -27, -76, -40, k = 90).

**Table 1.** Activations for Mentalizing Effects during the Vignette Period. Upper panel: whole brain analysis of the main effect of mentalizing during the vignette period at an FWE-corrected extent threshold of *p* < 0.05, initial cluster-forming height threshold *p* < 0.001 with a minimum cluster extend of 122 voxels. Lower panel: the reverse contrast for higher BOLD signal in the outcome compared to the belief condition at an FWE-corrected extent threshold of *p* < 0.05, initial cluster-forming height threshold *p* < 0.001 with a minimum cluster extend of 70 voxels.

| Structure | L/R | Cluster Size | x | y | z | Peak t |
| --- | --- | --- | --- | --- | --- | --- |
| *Belief > Outcome (vignette)* |  |  |  |  |  |  |
| Superior Temporal Gyrus | R | 554 | 48 | -28 | -1 | 6.89 |
| Temporal Parietal Junction (TPJ) | L | 497 | -48 | -61 | 26 | 6.62 |
| TPJ / Superior Temporal Gyrus | R | 380 | 48 | -55 | 23 | 6.39 |
| Inferior Frontal Gyrus | L | 122 | -36 | 20 | -19 | 6.26 |
| Medial Prefrontal Cortex (mPFC) | L | 422 | -6 | 47 | 35 | 5.70 |
| Middle temporal gyrus | L | 292 | -54 | -4 | -16 | 5.53 |
| Precuneus | L | 125 | -6 | -55 | 29 | 5.30 |
| *Outcome > Belief (vignette)* |  |  |  |  |  |  |
| Inferior parietal lobule | L | 14022 | -48 | -37 | 41 | -9.37 |
| Inferior frontal gyrus | R | 110 | 54 | 5 | 17 | -5.91 |
| Midbrain / thalamus | L | 70 | -9 | -19 | -4 | -4.69 |

**Table 2.**
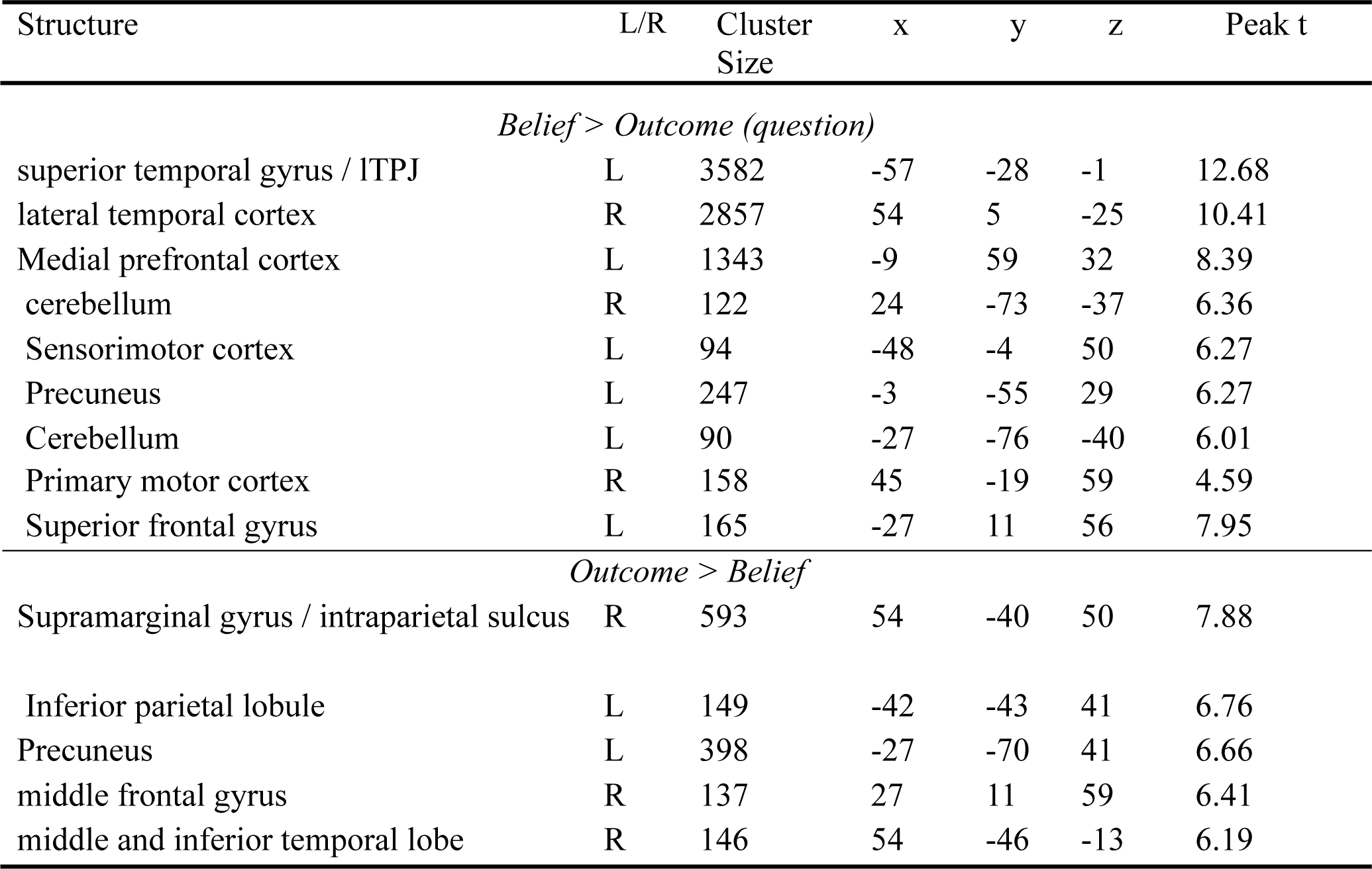
Activations for Mentalizing Effects during the Question Period. Upper panel: whole brain analysis of the main effect of mentalizing during the question period at an FWE-corrected extent threshold of *p* < 0.05, initial cluster-forming height threshold *p* < 0.001 with a minimum cluster extend of 90 voxels. The lower panel shows the reverse contrast reflecting higher BOLD signal in the outcome compared to the belief condition at an FWE-corrected extent threshold of *p* < 0.05, initial cluster-forming height threshold p < 0.001 with a minimum cluster extend of 137 voxels.

#### Effects of threat on activation patterns during vignette and question periods

The main goal of current study was to understand the impact of anxiety on the neural circuitry of social cognition. Our study enables us to investigate these effects on two separate social cognitive processes, namely belief formation, which occurs during the vignette period as subjects learn about the interactions between two partners, and belief inferences, which occurs in the question period as subjects apply what they have learned to answer targeted questions about characters’ beliefs. To identify the effects of anxiety on activation patterns during both the vignette and question periods we first examined the main effect of the threat manipulation. We performed two tests of this effect: given our previous results of suppressed mentalizing-related activity during trust decisions (Engelmann et al., 2019), we expected similar suppressive effects in the current task that were assessed via the contrast safe vs. threat. However, threat can also enhance task-specific activation, as previously demonstrated via enhanced negative subjective value coding in the insula during risky decision-making (Engelmann et al., 2015). We assessed this possibility by contrasting the threat to the safe condition.

During the vignette period, we identified increased threat-related activity in bilateral intraparietal sulcus (-27, -61, 35, k = 6522), which includes subclusters in ACC (18, 26, 35), dorsolateral PFC (-33, 14, 56), and cuneus / precuneus (27, -67, 35) (**Figure S4A**). Many of these regions overlap with canonical attention regions, as identified via a conjunction analysis between the current results and the neurosynth meta-analysis for attentional control (**see Figure S5A**). These results tentatively suggest that threat of shock lead to enhanced attention-related activation during the vignette period. This, however, is a non-specific effect that could indicate enhanced sustained attention to the vignette task, but also to the environment due to the possibility of receiving an electric shock throughout this prolonged period. No significant suppression was found during belief formation in the vignette period (**Table 3**).

**Table 3.** Activations table showing Threat Effects during the Vignette Period. Upper panel: whole brain analysis of the main effect of threat enhancement during the vignette period at an FWE-corrected extent threshold of p < 0.05, initial cluster-forming height threshold p < 0.001 with a minimum cluster extend of 138 voxels. Lower panel: no significant effect was found in the suppression contrast.

| Structure | L/R | Cluster size | x | y | z | Peak t |
| --- | --- | --- | --- | --- | --- | --- |
| *Main effect: Threat > Safe* |  |  |  |  |  |  |
| Intraparietal sulcus | L | 6522 | -27 | -61 | 35 | 7.27 |
| *corpus callosum (subcluster)* | R |  | 24 | 23 | 20 | 7.04 |
| *Anterior cingulate cortex (subcluster)* | R |  | 18 | 26 | 35 | 6.40 |
| *fusiform gyrus (subcluster)* | R |  | 24 | -64 | -10 | 5.99 |
| *V1 (primary visual cortex) (subcluster)* | R |  | -27 | -79 | 11 | 5.99 |
| *parahippocampal gyrus (subcluster)* | R |  | 33 | -34 | 23 | 5.97 |
| *DLPFC (subcluster)* | L |  | -33 | 14 | 56 | 5.92 |
| *parietal lobule (subcluster)* | L |  | -24 | -76 | 41 | 5.89 |
| *fusiform gyrus (subcluster)* | L |  | -38 | -67 | -10 | 5.81 |
| *middle occipital lobule* | L |  | -33 | -78 | 0 | 5.73 |
| *premotor cortex / frontal eye field* | L |  | -42 | -7 | 53 | 5.67 |
| *angular gyrus* | R |  | 42 | -64 | 35 | 5.65 |
| *corpus callosum (subcluster)* | L |  | -21 | 28 | 23 | 5.63 |
| *Cuneus / precuneus (subcluster)* | R |  | 27 | -67 | 35 | 5.56 |
| *Intraparietal sulcus (subcluster)* | R |  | 33 | -70 | 35 | 5.47 |
| R. Middle / Superior frontal gyrus | R | 138 | 36 | 5 | 53 | 5.33 |
| *Main effect: Safe > Threat* |  |  |  |  |  |  |
| No significant activations |  |  |  |  |  |  |

During the question period, we identified significant suppression due to threat in bilateral TPJ (-63, -52, 38, k = 151; 54, -46, 53, k = 139), bilateral inferior frontal gyrus (-42, 20, 5, k =79; 54, 29, -7, k = 64) and left putamen (-15, -13, 2, k = 330) (**Figure S4B**). These results are consistent with the notion that one effect of threat is to suppress specifically those regions that are importantly involved in solving ongoing task demands, which in the current task are regions involved in taking the perspective of others. No enhanced activity was found during belief inferences in the question period (**Table 4**).

**Table 4.** Activations for Threat Effects during the Question Period. Upper panel: whole brain analysis of the main effect of threat suppression during the question period at an FWE-corrected extent threshold of *p* < 0.05, initial cluster-forming height threshold *p* < 0.001 with a minimum cluster extend of 64 voxels. Lower panel: no significant effect was found in the enhancement contrast.

| Structure | L/R | Cluster size | x | y | z | Peak t |
| --- | --- | --- | --- | --- | --- | --- |
| *Main effect: Safe > Threat* |  |  |  |  |  |  |
| L. supramarginal gyrus / TPJ | L | 151 | -63 | -52 | 38 | 5.67 |
| L. inferior frontal gyrus | L | 79 | -42 | 20 | 5 | 5.07 |
| L. thalamus / putamen | L | 330 | -15 | -13 | 2 | 4.93 |
| R. supramarginal gyrus / TPJ | R | 139 | 54 | -46 | 53 | 4.87 |
| R. inferior frontal gyrus | R | 64 | 54 | 29 | -7 | 4.60 |
| *Main effect: Threat > Safe* |  |  |  |  |  |  |
| No significant activations |  |  |  |  |  |  |

#### Effects of anxiety on activation reflecting belief-formation during the vignette and belief inferences during the question period

To assess to what degree the effects of threat on neural activations during belief formation and belief inferences affected specifically social cognition regions involved in mentalizing, we first probed our fMRI data for the interaction between the factors Threat and Belief. We do not find significant interaction effects during either period, reflecting that anxiety did not *specifically* impact activity related to mentalizing. To probe our data for non-specific effects of anxiety, we performed conjunction analyses that assess whether the threat-related enhancements during the vignette period and suppressions during the question period occur in social cognition regions involved in mentalizing in the current task. We therefore performed conjunction analyses by multiplying the FWE corrected maps (cluster-forming threshold of *p* < 0.001 with FWE cluster-level correction) for the main effects of mentalizing and threat from the previous analyses.

In the vignette period, we find significant overlap between regions that are showing *enhanced* activity due to threat and regions involved in belief-formation. These regions include the left medial temporoparietal junction (-42, -61, 26, k = 27) and the precuneus (-3, -55, 32, k = 45). **Figure 4 (and Table 5**) shows these regions together with the time courses for the two main effects that simultaneously show enhanced belief-based activation as well as enhanced threat related activation in these regions.

**Figure 4.**
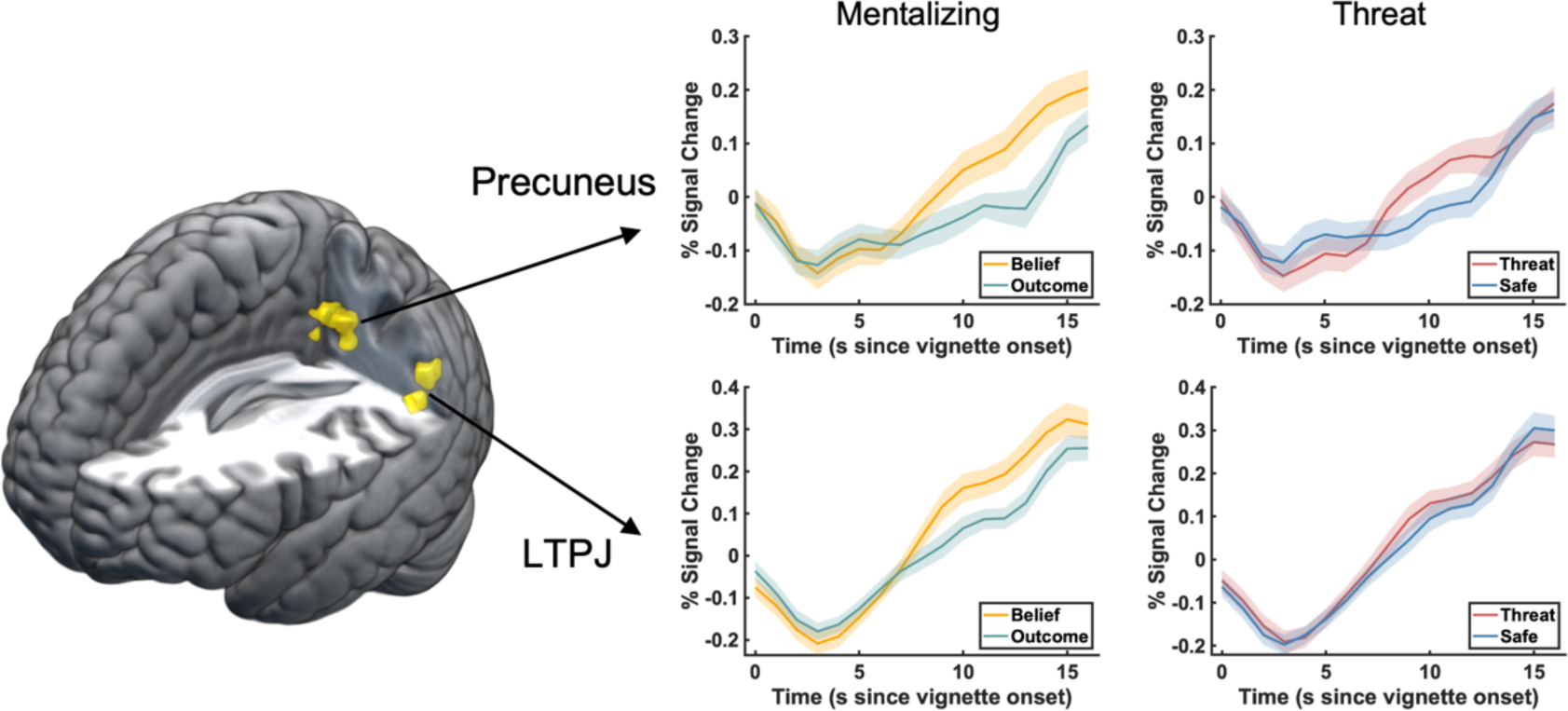
Conjunction analysis of activation reflecting belief formation and threat-enhancement during vignette period. Areas are shown that activate during mentalizing (belief > outcome) and, at the same time show enhancement by threat (threat > safe) in the vignette period. Two regions were identified in this conjunction analysis including medial left TPJ and precuneus (also see **Table 5**). Inlets show time courses of significant activations plotted separately for the mentalizing and threat main effects. Time courses were extracted from all voxels in the regions identified by conjunction analyses.

**Table 5.** Conjunction analysis of mentalizing and threat manipulations during vignette and question period respectively. Results from conjunction analyses identifying regions simultaneously showing (1) enhanced mentalizing-related and enhanced threat-related activity and during the vignette period, and (2) enhanced mentalizing-related and suppressed threat-related activity in the question period. Individual maps that were entered into conjunction analyses were thresholded at cluster level with FWE corrected alpha < 0.05. Conjunction analyses were conducted under the conjunction null hypothesis, requiring that all comparisons in the conjunction are individually significant (see Nichols et al., 2005).

| Structure | L/R | Cluster Size | x | y | z |
| --- | --- | --- | --- | --- | --- |
| *Conjunction of mentalizing effect and the enhancement of threat during vignette period* |  |  |  |  |  |
| Superior Temporal Gyrus | L | 27 | -42 | -61 | 26 |
| Precuneus | L/R | 45 | -3 | -55 | 32 |
| *Conjunction of mentalizing effect and the suppression of threat during question period* |  |  |  |  |  |
| Inferior frontal gyrus | L | 29 | -54 | 23 | 5 |
| Putamen | L | 8 | -28 | 8 | 5 |
| Temporoparietalparietal junction | L | 83 | -60 | -61 | 20 |
| Inferior frontal gyrus | R | 28 | 51 | 26 | -7 |
| Temporoparietalparietal junction | R | 22 | 60 | -58 | 35 |

In the question period, we find significant overlap between regions that are involved in mentalizing and regions that are significantly *suppressed* by threat. The conjunction map is shown in **Figure 5** (also see **Table 5**) and reveals overlap in bilateral TPJ (left: -60, -61, 20, k = 83; right: 60, -58, 35, k = 22), bilateral IFG (left: -54, 23, 5, k = 29; right: 51, 26, - 7, k = 28), and also left putamen (-28, 8, 5, k = 8). Time courses were extracted from clusters that show significant conjunction and are plotted separately for the mentalizing and threat main effects.

**Figure 5.**
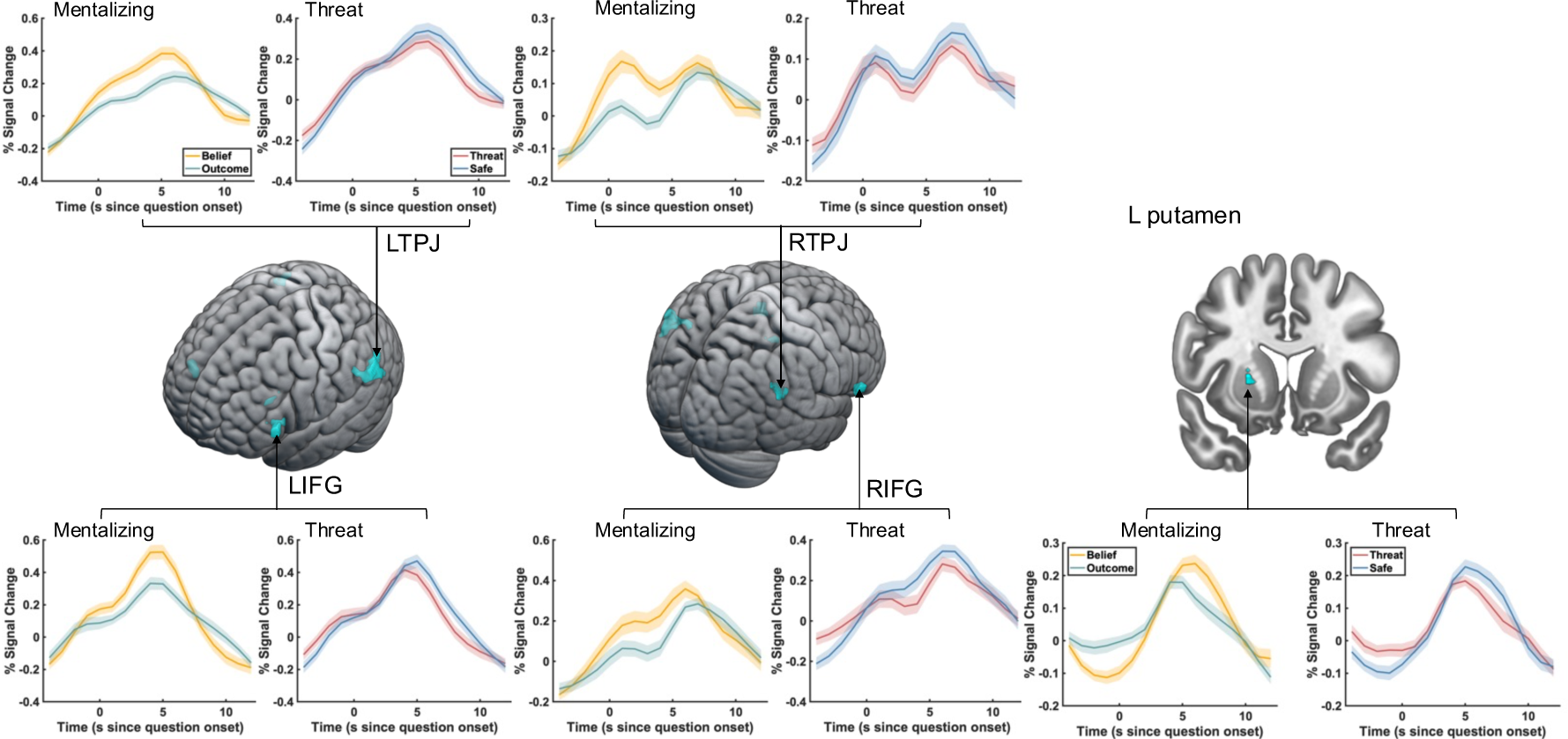
Conjunction analysis of activation reflecting belief inferences and threat-suppression during question period. Areas are shown that activate during mentalizing (belief > outcome) and, at the same time show suppression by threat (safe > threat) in the question period. Five main clusters were identified in this conjunction analysis including bilateral IFG, bilateral TPJ, and left putamen (also see **Table 5**). Inlets show time courses of significant activations plotted separately for the mentalizing and threat main effects. Time courses were extracted from all voxels in the regions as identified by conjunction analyses.

### Functional Connectivity Results using General PsychoPhysiological Interaction (gPPI) Analyses

To identify the effects of anxiety on connectivity between regions showing activity that reflects both mentalizing and threat-related changes in activity, we conducted generalized PPI analyses using as seeds the regions from the conjunction analyses reported in **Figure 4** for the vignette period and in **Figure 5** for the question period. We first tested the main effect of threat on connectivity during the vignette and question periods, to identify the effects of threat on connectivity while our participants were generating beliefs (during the vignette period) and making belief-based inferences (during the question period). During the vignette period, the main effects of threat identified suppressed connectivity within the social cognition network. As shown in **Figure 6A**, we found threat-induced suppression in connectivity between the seed in precuneus and its targets in right TPJ (58, -52, 26, k = 243), as well as a posterior region of dmPFC (16, 30, 50, k = 138). We did not find any *increased* connectivity as a function of threat during belief formation. During the question period, shown in **Figure 6B**, threat suppressed the connectivity between the right IFG seed and its target in left supramarginal gyrus (-56, -42, 8, k = 247), and increased connectivity between the left TPJ seed and its target in posterior precuneus (6, -82, 38, k = 136).

**Figure 6.**
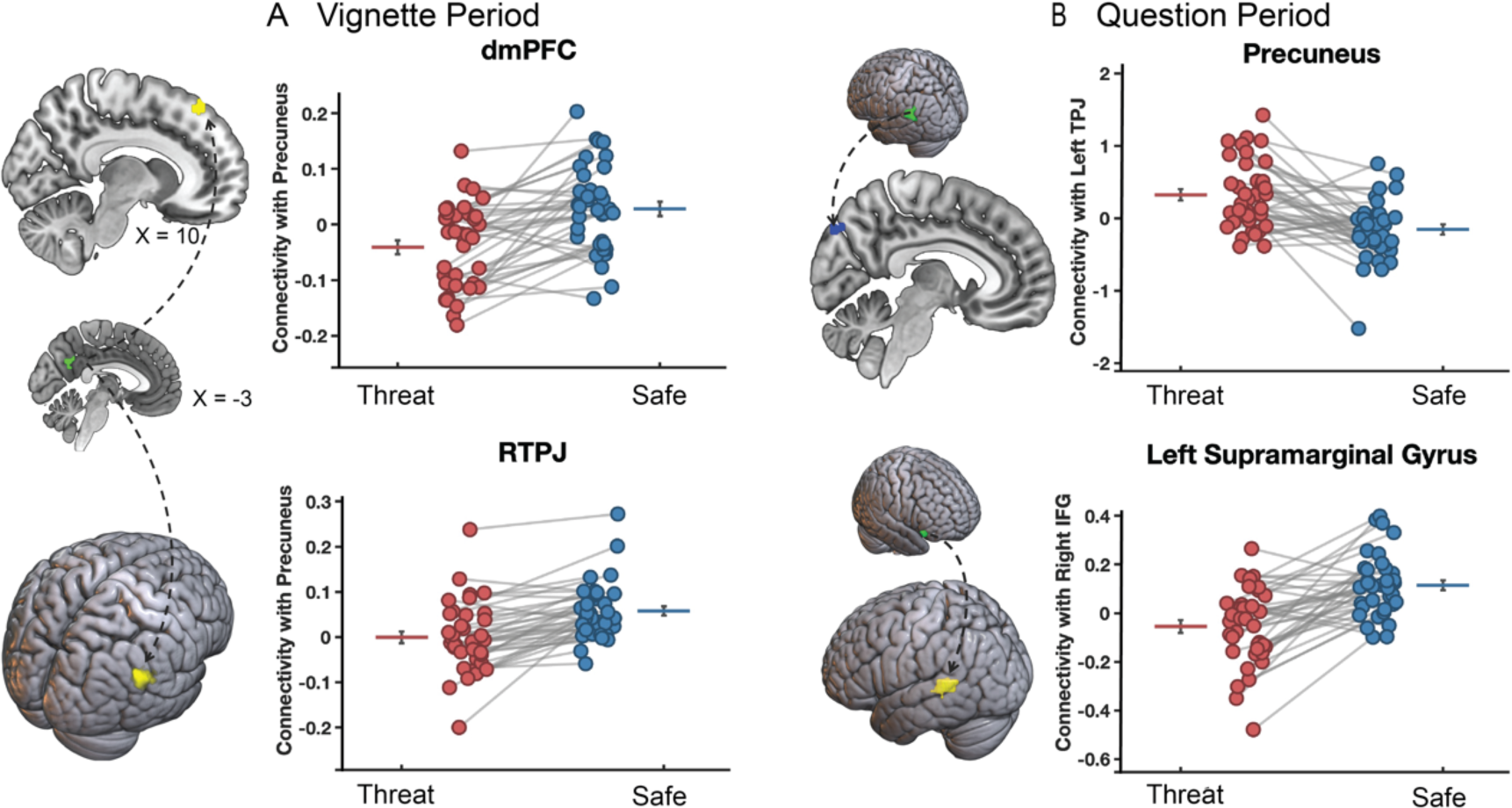
Main effect of threat on connectivity during vignette (A) and question (B) periods. (A) During the vignette period, the connectivity between the precuneus seed region and dmPFC and rTPJ target regions is significantly suppressed by threat. (B) During the question period, the connectivity between left TPJ and precuneus is enhanced, while a suppression is found between right IFG and left SMG.

### Specific suppression of belief-based activity

After identifying the main effect of threat on connectivity, we next conducted follow-up analyses probing for *specific* effects of Threat on belief-based connectivity compared to outcome-based connectivity via an interaction analysis. During the vignette period we were not able to identify such specific effects. However, during belief inferences in the question period, we found significant threat-related suppression of connectivity in the belief but not the outcome condition between the left putamen seed and its targets in bilateral intraparietal sulcus (IPS; right: 24, -52, 38, k = 139; left: -32, -60, 38, k = 168) and left dlPFC (-42, 18, 44, k = 334), but not during outcome-related inferences (**Figure 7**). These regions fall within the canonical attention network (see conjunction analysis in **Figure S5B**), which tentatively suggests reduced connectivity with attentional control regions when participants made inferences about others’ beliefs under conditions of anxiety.

**Figure 7.**
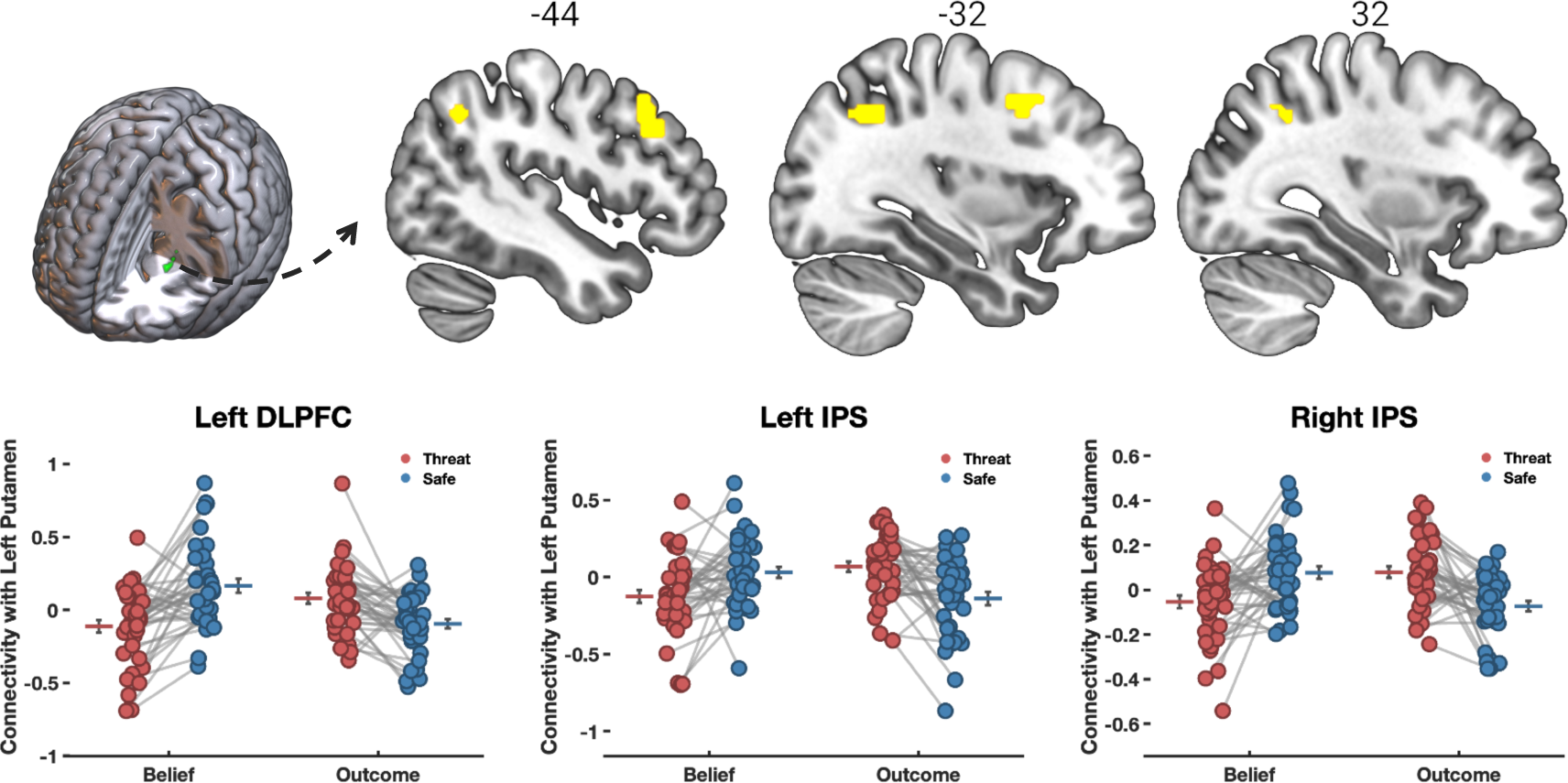
Threat specifically suppresses connectivity between left putamen and its targets in IPS and dlPFC. The figure shows significant threat-related suppression of belief-based connectivity during the question period, reflected by the interaction between the threat and belief factors. Connectivity between the putamen seed region and its targets in IPS and dlPFC is specifically suppressed in the Belief condition, but not in the outcome condition within this network.

Connectivity-behavior analyses further confirm the connection between regions involved in both mentalizing and threat suppression during the question period and their suppressed effective connectivity with attentional control regions. Specifically, we inspected the relationship between threat-related connectivity (threat main effect contrast: safe > threat) and dispositional distress. Dispositional distress is a latent variable obtained from factor analyses of multiple questionnaires measured before the start of the experiment. Distress has high loadings (> 0.7) on the State-Trait Anxiety Inventory (STAI), NEO5 neuroticism, Beck’s Depression Inventory (BDI), and Perceived Stress Scale (PSS) (see the factor analysis results reported in the supplement in **Table S3** and **Figure S3**). We found that dispositional distress significantly modulated threat-related suppression of connectivity between the left TPJ seed and left IPS, as well as between the right IFG seed and left sensorimotor cortex. These results indicate that left TPJ – IPS (and IFG-PCG) connectivity is suppressed to a larger extent for subjects reporting significant trait distress.

**Figure 8.**
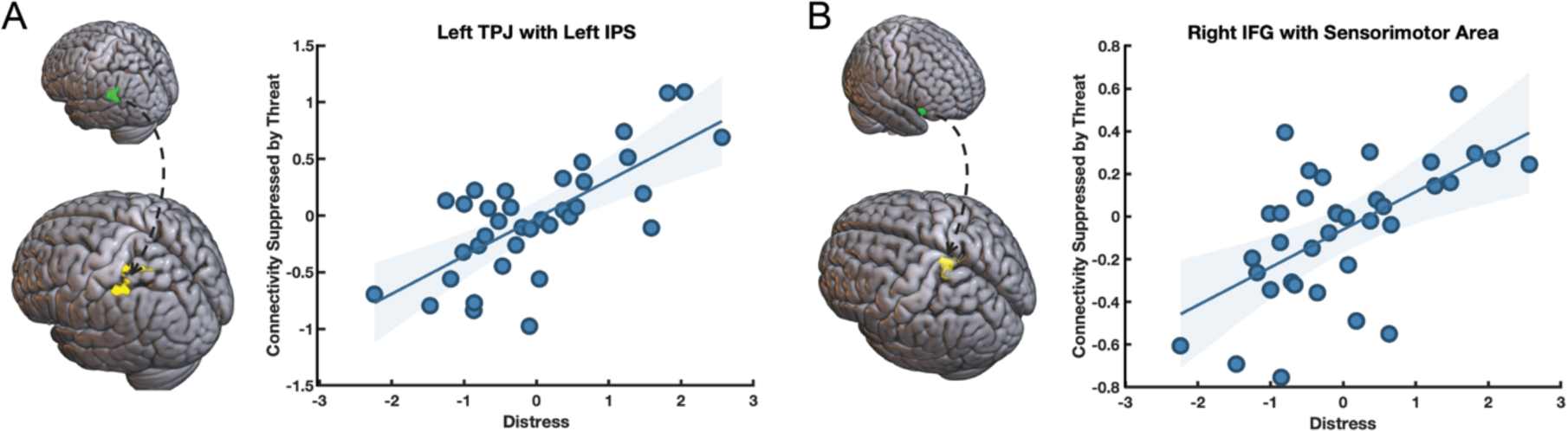
Suppression of connectivity between left TPJ and left IPS and right IFG and left sensorimotor cortex is moderated by dispositional distress. (A) the seed in left TPJ (green) shows suppressed connectivity with its target in left IPS (yellow) as a function of dispositional distress. (B) The seed in right IFG (green) shows suppressed connectivity with its target in left sensorimotor cortex (yellow). The y-axis reports the main effect of threat (safe - threat), such that reduced connectivity under threat compared to safety leads to greater contrast values reflecting enhanced threat-suppression.

## Discussion

The current experiment invesitgated the impact of anxiety on the neural signature of mentalizing by asking participants to complete two versions of the false-belief task, an economic games and a standard version, under the threat of electrical shock. We observed a clear effect of threat on arousal and self-reported emotion, but found only marginal effects of threat on behavior reflected by slightly improved average performance under threat. Similar to our previous study (Engelmann et al., 2019), the behavioral effects of anxiety were not specific to mentalizing, but also extended to the outcome control condition as indicated by an absence of a significant interaction between threat and belief. This has the advantage that, on average, the two threat conditions are well matched in performance leading to little performance-based (RT and accuracy) distortions of the fMRI signal in our analyses of the effects of threat on the neural correlates of mentalizing (see also Engelmann et al., 2019; Hein et al., 2016; Wilkinson & Halligan, 2004).

Our fMRI results replicate and extend prior work on the neural signature of mentalizing. First, we replicate prior results showing that mentalizing is associated with activity in areas within the social cognition network including bilateral temporal pole, right superior temporal gyrus, bilateral TPJ, precuneus / PCC, and also mPFC (see **Figure S3** and our previous analysis of this data reported in Chang et al., 2023). This shows that in analyses that control for threat, average activation patterns related to mentalizing follow previously reported patterns, showing activity during belief formation (vignette period) and belief inferences (question period) in a core social cognition network. Importantly, we extend prior work by demonstrating the effects of anxiety on the neural correlates of belief formation and belief inferences. When contrasting threat of shock to safety, our results demonstrate (1) enhanced activity in medial-TPJ and precuneus under threat during belief formation in the vignette period, and (2) suppression of activity in a wider social cognition network during belief inferences in the question period. The latter finding suggests that regions that are involved in taking the perspective of others are the same regions that show suppression under conditions of threat and anxiety. We tested this via a conjunction analysis between the contrast Belief vs. Outcome, reflecting mentalizing, and the contrast Safe vs. Threat, reflecting threat suppression. We found significant overlap between mentalizing and threat suppression in bilateral TPJ, bilateral IFG, and left putamen (**Figure 5**). This finding is consistent with our hypothesis that anxiety suppresses the neural underpinnings of mentalizing. However, our results indicate that the suppressive effects of anxiety are non-specific given the absence of significant interactions between Threat and Belief in our behavioral and fMRI models. In line with our previous fMRI study investigating the impact of anxiety on the neural correlates of trust decisions (Engelmann et al., 2019), we show here that anxiety also suppressed TPJ and IFG when participants made inferences about others’ beliefs in two iterations of the false belief task, a standard and economic-games version, both of which require mentalizing. These results therefore lend further support to the notion that one effect of anxiety is to suppress activity in social cognition regions when current task demands draw on mentalizing abilities. Thus, in the context of both economic games and false belief tasks that rely on mentalizing abilities, anxiety can suppress the activity and connectivity of social cognition regions (Engelmann et al., 2019).

The finding that bilateral TPJ is suppressed in the presence of threat confirms and extends previous results demonstrating suppressed TPJ activity after social stress induction during RMET performance (Nolte et al., 2013) and suppressed activity and connectivity during active intersocial behavior that require mentalizing, i.e., trust decisions, under threat of shock (Engelmann et al., 2019). The TPJ has been repeatedly implicated in mentalizing and theory of mind (Decety & Lamm, 2007; Mar, 2011; Mitchell, 2009; Molenberghs et al., 2016; Schurz et al., 2014; Van Overwalle, 2009) and in the current study we replicate this result in the context of a standard false belief task and a recent extension of this task that describes bilateral interactions in economic games (for a more in-depth discussion of the novel economic-games false belief task and its neural correlates, see Chang et al., 2023). Importantly, we find that the TPJ simultaneously shows a main effect of mentalizing and of threat, particularly during belief inferences. Moreover, the activation clusters of these main effects overlap spatially as identified via conjunction analyses. Jointly, these results suggest threat-related, but non-specific suppression of activity in the TPJ during belief inferences.

The finding of suppression in IFG under conditions of threat is consistent with its role in cognitive control and emotion regulation (Aron et al., 2014; Bari & Robbins, 2013; Hampshire et al., 2010). This region is also a central part of an extended social cognition network (e.g., Arioli et al., 2021) and likely plays a role in mentalizing by inhibiting self-referential thought to enable taking the perspectives of others. This antagonistic connection between self-referential thought and mentalizing has been confirmed by a number of prior behavioral, fMRI and lesion studies. Schneider et al. (2012) used a dual-task setup consisting of an auditory N-back task and a simultaneous movie where an actor held a false belief about an object’s location. Under high cognitive load, participants’ eye movements no longer reflected the actor’s false belief, indicating that disrupting executive functions via cognitive load can inhibit mentalizing. These behavioral results are likely linked to vlPFC function. A previous fMRI study using both a ToM and Stop Signal Task in the same subjects revealed that bilateral IFG is involved both in mentalizing and motor inhibition (Van der Meer et al., 2011), lending further support to the notion that IFG is important for both inhibitory and social cognitive processes. Likewise, the role of IFG in inhibiting self-referential thought was also demonstrated in a neuropsychological study which indicated that patients with VLPFC lesions made egocentric errors in a social cognition task (Samson et al., 2005). Finally, IFG activity has been shown to be suppressed in the context of anxiety and stress. One study demonstrated that an anxious attachment style is associated with inhibited bilateral IFG activity during mentalizing, compared to avoidant attachment styles (Schneider-Hassloff et al., 2015). Decreased VLPFC activity has been demonstrated in a working memory task under conditions of stress. Specifically, Koric et al. (2012) utilized two working memory tasks with varying levels of performance-based stress. They found that in healthy subjects, right VLPFC activity was suppressed in the stressful condition compared to the less stressful condition. In the same vein, our conjunction analysis showing simultaneous main effects of mentalizing and threat suggests that anxiety can disrupt mentalizing processes by suppressing cognitive control regions such as IFG.

### Connectivity results

Our functional connectivity analyses showed largely suppressive effects of anxiety, further supporting the suppressive effects of threat on activity in social cognition regions. We found threat-based suppression of connectivity both during the vignette period, specifically between the precuneus seed region and target regions in dmPFC and rTPJ, as we all as during the question period, where the connectivity between right IFG and left SMG was suppressed. These main effects reflect that connectivity is largely suppressed by threat during vignette and question periods. One exception is the connectivity between left TPJ and posterior precuneus, which was enhanced by threat during the question period. Such non-specific effects of threat could indicate enhanced effort in sustaining attention to the task at hand in the presence of threat, but also enhanced attention to a threatening environment due to the possibility of receiving an unpredictable electric shock throughout a prolonged period. The patterns of connectivity observed between the precuneus and other nodes of the social cognition network in our study aligns with the notion that the precuneus may be functionally heterogenous (Willbrand et al., 2022). This concept resonates with previous research that reported a dissociation between the ventral and dorsal parts of the PCC (Leech et al., 2011). Specifically, Leech et al. (2011) demonstrated that connectivity between the ventral-PCC and default mode network (DMN) nodes was suppressed under high-cognitive load conditions, whereas dorsal-PCC showed increased connectivity with the DMN in the same condition.

In addition to these non-specific effects of threat, we also found specific threat-related suppression during mentalizing in the question period, i.e., connectivity that is specifically suppressed in the belief relative to the outcome condition. Such interaction effects were identified for the connectivity between the left putamen seed and its targets in IPS and dlPFC. They reflect specific threat-based suppression of connectivity during mentalizing, but not during reasoning about events that happen to another person and economic game payouts during the outcome condition. Our functional connectivity results therefore demonstrate an interaction between the reward system, represented by the putamen seed, and frontal-parietal attentional systems, represented by IPS and dlPFC, when participants engage in social cognition under threat (Starr et al., 2011). In a conjunction analysis reported in **Figure S5**, we show that these target regions overlap with attentional control regions as identified by neurosynth meta-analyses. These results point to the possibility that social cognitive processes receive support from additional networks, particularly attentional control regions, and that these connections are suppressed during threat. This notion is further supported by behavior-connectivity analyses, which showed that connectivity between left TPJ and left IPS is moderated by dispositional distress. Specifically, during threat, connectivity is suppressed to a larger extent for subjects showing higher levels of dispositional distress. Jointly, these results indicate an interesting interaction between mentalizing and attentional regions that is moderated by the presence of threat and the dispositional distress of a person.

One interpretation of the current results of reduced interconnectivity between social cognition and attentional control regions rests on results commonly reported for anxiety disorder, namely threat-related attentional biases, such as difficulties in disengaging attention from threatening and invalid cues (e.g., Pacheco-Unguetti et al., 2011; Yiend & Mathews, 2001). Meta-analyses show that such attentional biases are reliably found in anxious individuals (Bar-Haim et al., 2007; Cisler & Koster, 2010). Given that attentional networks are disrupted in anxiety disorders (Sylvester et al., 2012; Williams, 2016), and that such disruption is associated with the extent of dispositional anxiety (Forster et al., 2015), the reduced connectivity between these regions could reflect reduced flexibility in social cognitive processes under conditions of anxiety. These effects may be driven by enhanced attention to the threatening context, which limits the cognitive resources that can be dedicated to the task required of participants (see also Engelmann et al., 2015; Engelmann et al., 2017). However, our discussion of the connectivity results is rather speculative at this point and future research is needed to identify the specific interactions between social cognitive and attentional networks. One important task will be to identify how connectivity relates to behavioral changes, which we were not able to identify in the current study due to generally high-performance levels.

### Limitations

Despite careful experimental design and well-matched control conditions, a number of potential limitations are worth mentioning. First, while we calibrated the task in two pilot studies (reported in Chang et al., 2023), performance in the scanning environment improved and was close to ceiling level. The false belief task used here was therefore not optimal for revealing the behavioral effects of threat. Future studies on the effects of threat on mentalizing performance should therefore consider a more difficult task to maximize their ability to detect the behavioral effects of threat. This could be accomplished by redesigning the questions from the current task (for instance by including additional answer options), or using a more difficult task such as reasoning in a betting game scenario (Carpenter & Robbett, 2022). On the flip side, however, current results are consistent with previous observations that experimental treatments, while having similar effects on behavior, can differentially impact mental operations and the activation patterns associated with those (Engelmann et al., 2019; 2015; Hein et al., 2016, Wilkinson and Halligan, 2004). Moreover, since the purpose of the current study was to understand the effects of threat in terms of brain activity and connectivity, it can be desirable that there are no differences in behavioral responses across conditions as this reduces performance-based distortions of the fMRI signal (Engelmann et al., 2015; Engelmann et al., 2019; Hein et al., 2016; Wilkinson & Halligan, 2004). Furthermore, it should be mentioned that the outcome condition in the economic-games context is associated with slightly reduced accuracy. This condition, in which participants were asked to compute the payoffs for played involved in trust and ultimatum game interactions, served as an important control condition for the economic-games false belief task. Our approach in the current paper was to collapse across the two version of the false-belief task we included here to enhance statistical power, as we show in a previous paper that these two tasks lead to very similar activation patterns within the social cognition network (for detailed discussion see Chang et al., 2023). Moreover, we control for this condition in our analyses via the inclusion of the Task Domain regressor in all behavioral and fMRI models.

A further limitation of our study is the lack of jitter between the vignette and question periods of the task. Jitter would have allowed for better separation of the hemodynamic response across these periods. Despite this, our decision not to include jitter was based on several factors: (i) to facilitate comparison with previous studies (e.g., Saxe & Kanwisher, 2003; Young et al., 2010a; Liane Young et al., 2010b), (ii) to reduce the cognitive burden on participants, and (iii) to keep the experiment duration reasonable. Furthermore, this limitation is mitigated by the BOLD patterns observed in Figures 4 and 5, which show the expected responses during both task periods. Specifically, during the vignette period, BOLD responses peak at about 15s, reflecting sustained social cognitive processes, while during the question period, they peak at around 5s, reflecting transient social cognitive processes consistent with the average response time during that period. We show similar time courses in our previous analyses of the social-cognitive aspects of this dataset (see Chang et al., 2023), further mitigating this concern.

### Conclusion

Anxiety is a ubiquituous emotion that can have important adapative functions. Recent research has demonstrated that anxiety can have wide-ranging effects on cognitive processes that support ongoing behavior, including attention, memory and learning. This makes much sense, considering that one role of anxiety is to increase vigiliance and attention to avoid potentially adverse outcomes. Despite the extensive prior research on the impact of anxiety on cognition, we identified a gap in the neurobiological literature on the effects of anxiety on social cognitive processes. We address this gap in the current paper by inducing anxiety using the well-established threat-of-shock procedure and assessing its behavioral and neural impact on mentalizing. Mentalizing is an important social cognitive process that enables us to understand the beliefs and intentions of others and that is consistently associated with activity in a core social cognition network consisting of bilateral TPJ, dmPFC and precuneus. Our results suggest that incidental anxiety suppresses the activity and connectivity within the social cognition network during mentalizing, but does so in a non-specific manner. Our work further provides novel evidence for the involvement of the frontal-parietal attentional network during mentalizing, which shows threat-suppressed connectivity during mentalizing, but not during the control condition. These results underline the importance of large scale network interactions between social cognitive and attentional networks, further indicating that social cognitive processes are supported by attentional processes. One intriguing suggestion from our results is that reward system in general, and the putamen in this particular case, may serve as a hub enabling cross-talk between these two large-scale systems (see also Pessoa & Engelmann, 2010). Collectively, these results contribute to bridging the existing gap in the research on the neural effects of anxiety on specifically social processes, elucidating how incidental anxiety and dispositional anxiety modulate the neural circuitry underpinning social cognition. Our findings can inform future clinical research on anxiety disorders, such as SAD, OCD, and PTSD, which often show marked social deficits. By providing insights into the complex interplay between anxiety and its impact on multiple networks that support mentalizing, we hope to enable a way for developing more targeted and neurobiologically informed treatment approaches.

## Acknowledgements

We are grateful to Ava Q. Ma de Sousa and Fenke Paauwe for assistance with data collection. This work was supported by start-up funds from the Amsterdam School of Economics, which we gratefully acknowledge.

