## supplemental materials for "The impact of incidental anxiety on the neural signature of mentalizing"

### Supplementary Materials

#### Table S1. Behavioral Results from Mixed Effects Regressions

| | Accuracy | | | | Response Time (log) | | | |
|---|---|---|---|---|---|---|---|---|
| | Model 1 | | Model 2 | | Model 3 | | Model 4 | |
| | Chisq | Pr(>Chisq) | Chisq | Pr(>Chisq) | Chisq | Pr(>Chisq) | Chisq | Pr(>Chisq) |
| **Belief** | 15.404 | P<0.000 | 14.927 | P<0.000 | 4.469 | 0.035 | 4.485 | 0.034 |
| **Threat** | 2.363 | 0.124 | 1.857 | 0.173 | 0.422 | 0.516 | 0.406 | 0.523 |
| **Belief x Threat** | | | 0.163 | 0.686 | | | 0.356 | 0.551 |
| **Observations (ID)** | 3530 (37) | | 3530 (37) | | 3381 (37) | | 3381 (37) | |
| **AIC** | 1188.8 | | 1190.6 | | 3200.4 | | 3202.1 | |

ANOVA output from mixed linear and logistic regressions estimated respectively for accuracy and response times. All regressions include a Task Domain regressor of no interest. Below we report the maximal models that terminated without error in Wilkinson-Rogers notation for the main behavioral analyses reported in Table S1:

(1) Correct ~ Belief + Threat + Task Domain + (1 | Subject)
(2) Correct ~ Belief * Threat + Task Domain + (1 | Subject)
(3) log_rt ~ Belief + Threat + Task Domain + (1 + Belief | Subject)
(4) log_rt ~ Belief * Threat + Task Domain + (1 + Belief | Subject)

#### Table S2. Behavioral Results using the Balanced Integration Score (BIS) to control for potential speed-accuracy trade-offs

| | Model 1 | | Model 2 | |
|---|---|---|---|---|
| | Chisq | Pr(>Chisq) | Chisq | Pr(>Chisq) |
| **Belief** | 14.137 | P<0.000 | 14.140 | P<0.000 |
| **Threat** | 2.598 | 0.107 | 2.598 | 0.107 |
| **Belief X Threat** | | | 0.054 | 0.817 |
| **Observations (ID)** | 296 (37) | | 296 (37) | |
| **AIC** | 447.95 | | 449.900 | |

ANOVA output from mixed linear regressions estimated for BIS, which controls for speed-accuracy trade-offs across conditions. All regressions include a Task Domain regressor of no interest. Below we report the maximal models that terminated without error in Wilkinson-Rogers notation for the main behavioral analyses reported in Table S2:

(1) BIS ~ Belief + Threat + Task Domain + (1 + Task Domain | Subject)
(2) BIS ~ Belief * Threat + Task Domain + (1 + Task Domain | Subject)

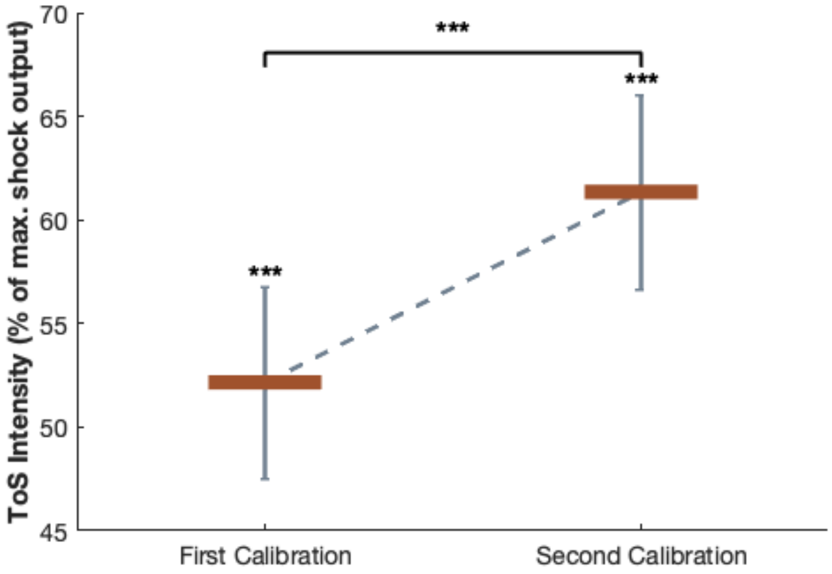


**Figure S1. Pain threshold calibration results.** The figure shows the outputs from the shock-intensity calibration staircase procedure after completion of the first and second calibrations. After the first calibration, the final calibration intensity was significantly different from the lowest intensity at 10% (average = 52.16%, $_{36}$ = 9.16, $p$ = $6.15\times10^{-11}$). The calibration output increased after the second calibration at the midway point of the experiment (average = 61.35%), as indicated by a paired t-test ($t_{36}$ = 4.62, $p$ = $4.81\times10^{-5}$).
$p < .1$ ~ ; $p < .05$ *; $p <.01$**; $p < .001$***

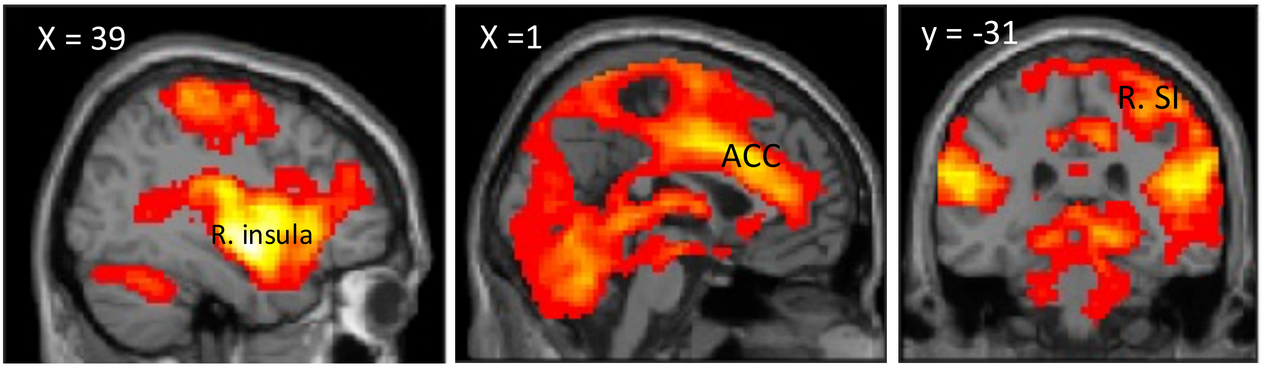


**Figure S2. Manipulation Check 3: Neural correlates during electrical shock administration within pain matrix.** The figure shows activation during shock administration. Activation within the pain matrix can be observed, including the insula, anterior cingulate cortex (ACC), and the right somatosensory cortex (R. SI) at n FWE-corrected extent threshold of $p < 0.05$ (39, -4, -10, k = 106, peak t = 16.64) and initial cluster-forming height threshold $p < .001$.

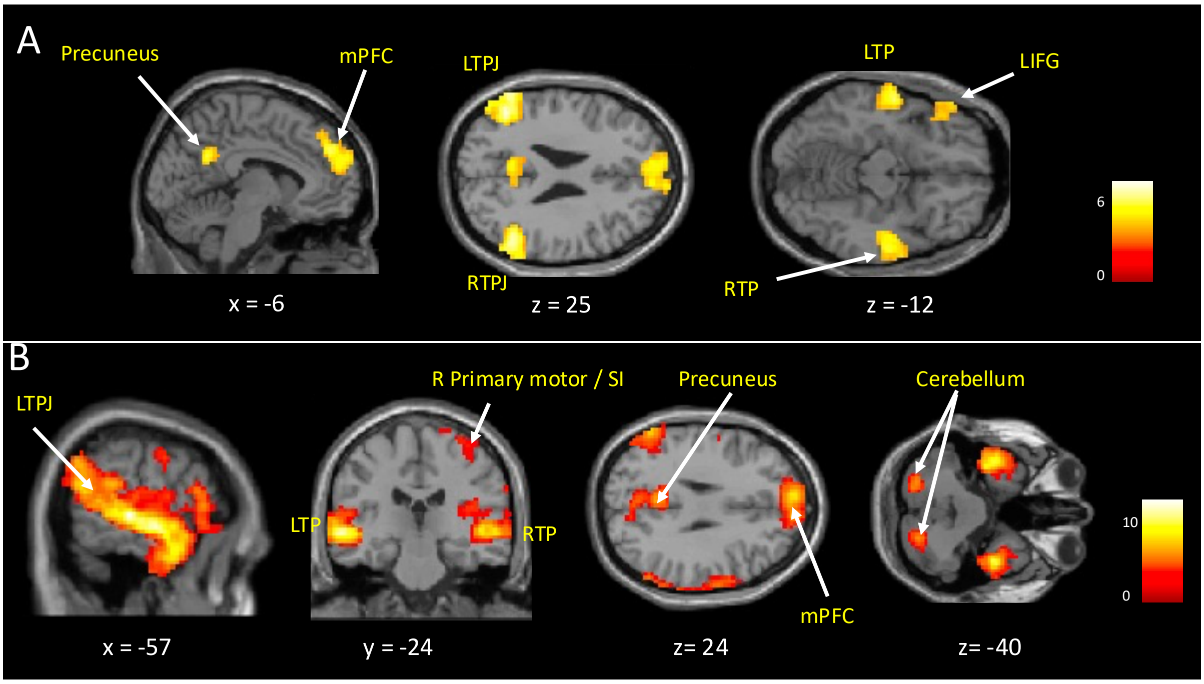


**Figure S3. FMRI results of mentalizing during belief formation and belief inferences in vignette and question periods respectively.** (A) Whole brain analysis of the main effect of mentalizing during the vignette period (FWE-corrected extent threshold of $p < 0.05$ (>= 122 voxels), initial cluster-forming height threshold $p < .001$). (B) Whole brain analysis of the main effect of mentalizing during question period (FWE-corrected extent threshold of $p < 0.05$ (>= 90 voxels), initial cluster-forming height threshold $p < 0.001$).

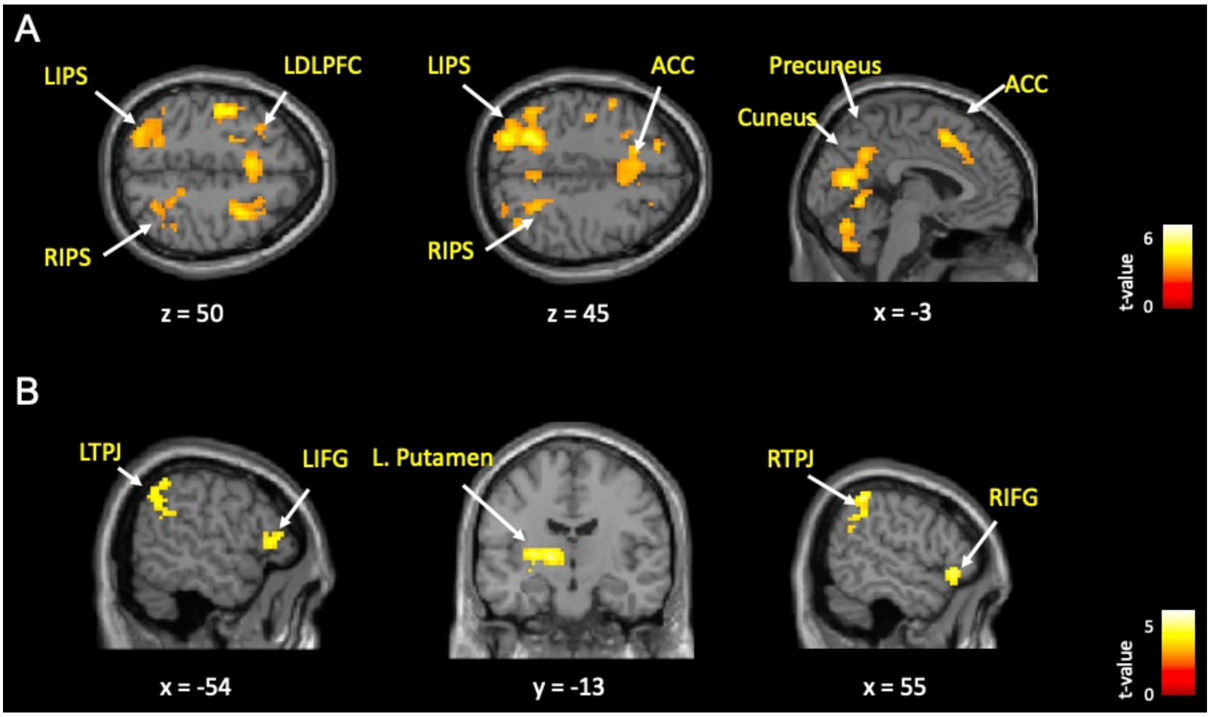


**Figure S4. FMRI results reflecting neural correlates of threat of shock.** (A) Whole brain analysis of main effect of threat (threat > safe) during vignette period (FWE-corrected extent threshold of $p < 0.05$ (>=138 voxels), initial cluster-forming height threshold $p < 0.001$). (B) Whole brain analysis of the reverse contrast during the question period (threat < safe) (FWE-corrected extent threshold of $p < 0.05$ (>= 64 voxles), initial cluster-forming height threshold $p < 0.001$).

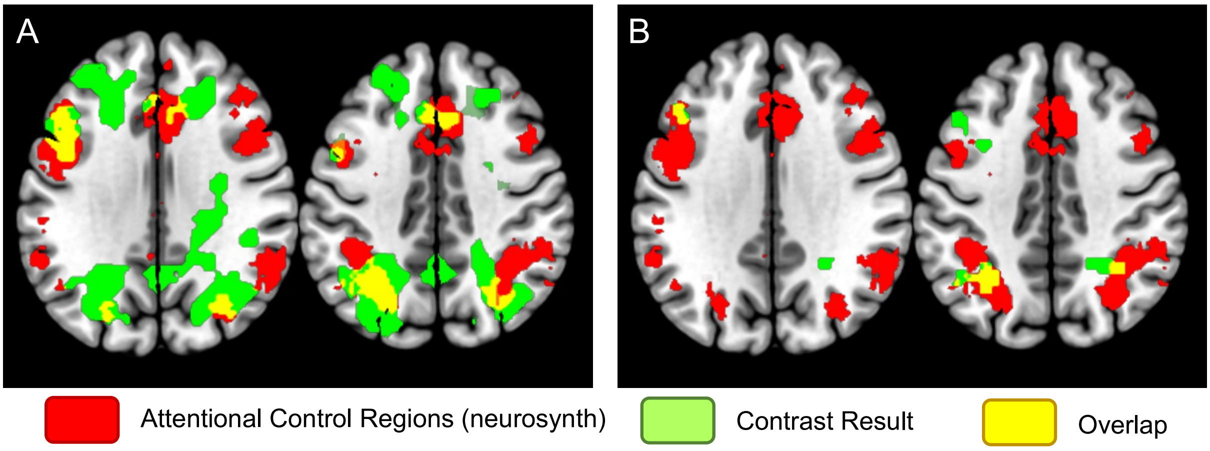


**Figure S5.** Conjunction analysis between attentional control regions (from neurosynth uniformity test "attentional control") and (A) regions showing significant enhancement of activity due to threat during the vignette period, and (B) target regions from putamen seed showing specific connectivity suppression by threat in the belief condition.

***Factor Analysis***

Participants completed a battery of personality tests adapted from our earlier study (Engelmann et al., 2019), which consists of the following measures: 1) Beck Depression Inventory-II (BDI-II) that measures individual's symptoms of depression; 2) Barrat Impulsiveness Scale (BIS-11) that assess individual's impulsiveness in scale of attentional impulsivity, motor impulsivity and cognitive impulsivity; 3) Behavioral avoidance/inhibition (BIS/BAS) scale that assess individual's appetitive motivations. While BIS (behavioral inhibition system) refers the avoidance of aversive event, BAS (behavioral approach system) refers the approach motives to desired event in subscale of drive, fun seeking, and reward responsiveness; 4) Experience in Close Relationships-revised (ECR-R-D) that measures the style of attachment anxiety and avoidance in adult intimacy and romantic relationship; 5) Interpersonal Reactivity Index (IRI) that measures individual's disposition of empathy and capability of theory of mind, consisting of sub-scale of perspective taking, fantasy scale, empathy concern, personal distress respectively; 6) MACH-IV test of Machiavellianism that measures individual's anti-social, manipulative and distrustful tendency; 7) Multidimensional Mood State Questionnaire (MDMQ) that assess individual's current mood, consisting three sub-scale of the extent of arousal and anxiety; 8) NEO Five-Factor Inventory (neuroticism, extraversion, openness to experience, agreeableness, conscientiousness); 9) Personal Norm of Reciprocity (PNR) measures individual's disposition of positive, negative and belief in reciprocity; Perceived Stress Scale (PSS) that measures the degree to which situations in participants' life are perceived as stressful; 10) Propensity to Trust Scale (PTS) that measures individual's propensity to trust others as well as their own character of trustworthy per se; 11) Social Desirability Scale (SDS), that assesses individual's propensity to behave in a socially desirable manner; 11) Sensation Seeking Scale, Form V (SSSV) that indexes individual's sensation seeking in scale of taking adventure, disinhibition, experience seeking and the susceptibility of being bored; State-Trait Anger Expression Inventory (STAXI) that assess individual's intensity and frequency of angry feelings; 12) State-Trait Anxiety Inventory, Form Y2 (STAI-Y2) that assess individual's state and trait anxiety (but see Knowles & Olatunji, 2020 for the controversy regarding the specificity of anxiety measure by this questionnaire; 13) Domain-Specific Risk-Taking (Adult) Scale that measures individual's risk attitude and the willingness to engage in risky activities. To maximize our subject-to-item ratio (to 3.56), we included the data from all subjects who completed this part of the study both in the context of the behavioral pilot (reported in Chang et al., 2023) and the fMRI experiment, and regardless of whether they did show up in the lab or not.

We performed an exploratory factor analysis using maximum likelihood estimation with orthogonal varimax rotation via the *factoran* function (Boedigheimer 2016) in MATLAB. As can be seen in the scree plot below (**Figure S3**), four factors were found to fall above the 5% cutoff value. **Table S3** shows the factor loadings of the subscores making up each factor. Specifically, we identified four latent factors that included: 1) **distress** with high loadings on depression (BDI), attentional impulsivity (BIS-11), motivation to avoid aversive outcomes, both anxious and avoidant romantic attachment styles (ECR-R), tendency to transpose fantasy into fictitious characters in novels or movies, neuroticism (NEO), distrust, anxiety, state and trait anger; 2) **prosociality** with high loadings on perspective taking, empathy concerns, low Machiavellianism, agreeableness, low in internalized norm of negative reciprocity; 3) **impulsivity** with high loadings on all sub-scales of impulsivity (BIS-11; attention, motor, and non-planning), low conscientious (NEO), low in internalized norm of positive reciprocity, and untrustworthy; 4) **fun seeking** with high loadings on all behavioral activation sub-scales (motivation to follow one's goal, sensitivity to pleasant stimuli in the environment and the motive to actively navigate the novel reward), and extraversion (NEO).

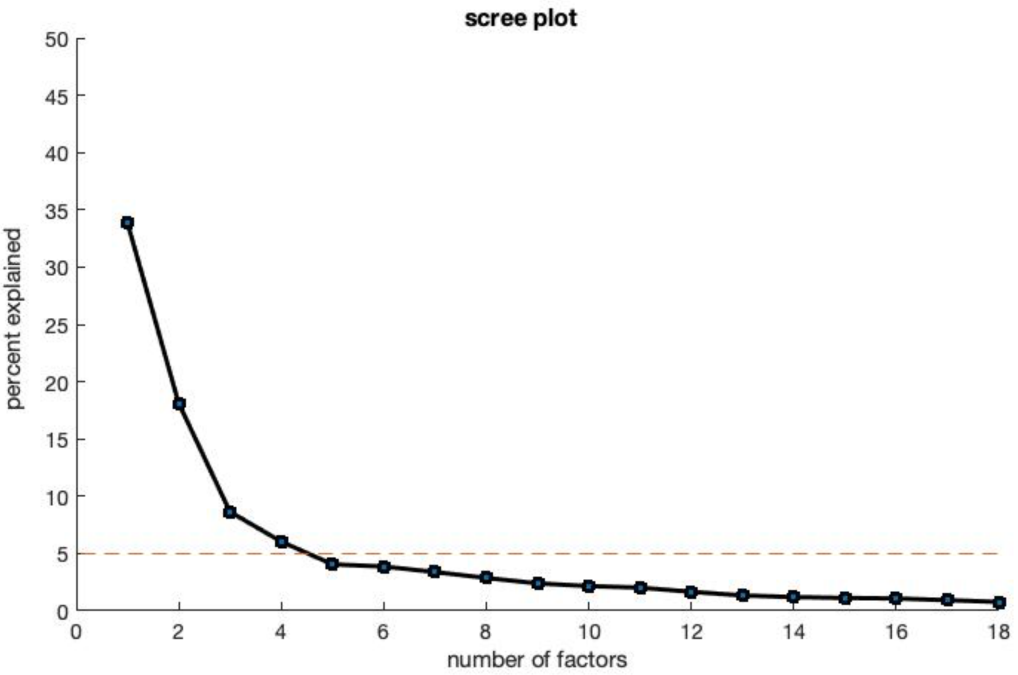


**Figure S3.** Scree plot of factor analysis for dispositional individual differences
Based on the scree plot results shown here, we retained four factors above that clearly fell above the 5% cutoff. To optimize the Factor Analysis, we tested different models by exclusing items that showed overall low loadings. The final model, which optimizes the Kaiser-Meyer-Olkin measure of sampling adequacy (0.77) and Cronbach (0.685) is reported in Table S3 and includes 34 items and four factors. The four factors include distress, prosociality, impulsivity, and fun seeking.

| Item | Distress | Prosociality | Impulsivity | Fun seeking |
|---|---|---|---|---|
| BDI | 0.72 | | | |
| BIS11 ATT | 0.53 | | 0.54 | |
| BIS11 MT | | | 0.58 | |
| BIS11 NP | | | 0.77 | |
| BA drive | | | | 0.58 |
| BA fun seeking | | | | 0.71 |
| BA reward response | | | | 0.57 |
| B Inhibit | 0.66 | | | |
| ECR anxious | 0.66 | | | |
| ECR avoidant | 0.42 | | | |
| IRI PT | | 0.61 | | |
| IRI FS | 0.42 | | | |
| IRI EC | | 0.56 | | |
| IRI PD | 0.57 | | | |
| MACH | | -0.67 | | |
| NEO5 neurotic | 0.88 | | | |
| NEO5 extravert | -0.35 | | | 0.66 |
| NEO5 openness | | 0.37 | | |
| NEO5 agreeable | | 0.86 | | |
| NEO5 conscientious | | | -0.77 | |
| PNR belief | | | | |
| PNR positive | | | -0.41 | |
| PNR negative | | -0.74 | | |
| PSS | 0.71 | | | |
| PTS trust | -0.56 | 0.53 | | |
| PTS trustworthy | | 0.34 | -0.51 | |
| SDS | | -0.39 | | |
| SSSV thrill | | | | 0.50 |
| SSSV EXP | | | | |
| SSSVdisinhibit | | | | 0.45 |
| SSSV bored | | | | |
| STAI | 0.89 | | | |
| STAXI state | 0.42 | -0.40 | | |
| STAXI trait | 0.39 | -0.58 | | |

**Table S3**. Factor loadings of Factor Analysis

Four suggested factors were identified, which can be referred to as the disposition of distress, prosociality, impulsivity, and fun seeking respectively. Kaiser-Meyer-Olkin measure of sampling adequacy is 0.77 and Cronbach is 0.685. Subject-to-item ratio: 3.56 (121 subjects vs. 34 items). Blanks are loadings below |0.35|.
